# P(*all-atom*) Is Unlocking New Path For Protein Design

**DOI:** 10.1101/2024.08.16.608235

**Authors:** Wei Qu, Jiawei Guan, Rui Ma, Ke Zhai, Weikun Wu, Haobo Wang

**Author notes:** Equal contribution. Correspondence to: Weikun Wu < >, Haobo Wang < >.

## Abstract

We introduce Pallatom, an innovative protein generation model capable of producing protein structures with all-atom coordinates. Pallatom directly learns and models the joint distribution P(*structure, seq*) by focusing on P(*all-atom*), effectively addressing the interdependence between sequence and structure in protein generation. To achieve this, we propose a novel network architecture specifically designed for all-atom protein generation. Our model employs a dual-track framework that tokenizes proteins into residue-level and atomic-level representations, integrating them through a multi-layer decoding process with “traversing” representations and recycling mechanism. We also introduce the `atom14` representation method, which unifies the description of unknown side-chain coordinates, ensuring high fidelity between the generated all-atom conformation and its physical structure. Experimental results demonstrate that Pallatom excels in key metrics of protein design, including designability, diversity, and novelty, showing significant improvements across the board. Our model not only enhances the accuracy of protein generation but also exhibits excellent sampling efficiency, paving the way for future applications in larger and more complex systems.

## 1. Introduction

The theoretical foundation of protein modeling has been built upon two key conditional probability distributions: P(*structure* | *seq*) and P(*seq* | *backbone*). The former, P(*structure* | *seq*), corresponds to the all-atom protein structure prediction task, which involves determining the three-dimensional structure of a protein given its amino acid sequence (Abramson et al., 2024; Jumper et al., 2021; Lin et al., 2023; Baek et al., 2023). The latter, P(*seq*| *backbone*), underpins the fixed-backbone design task, where the goal is to identify a sequence that will fold into a given protein backbone structure (Dauparas et al., 2022; Hsu et al., 2022). In summary, these probability distributions has successfully advanced the field of protein engineering.

With the advancement of deep learning in protein science, two distinct approaches for protein design have emerged. One approach is the protein hallucination (Anishchenko et al., 2021), which explores the landscape of a P(*structure* | *seq*) model using Monte Carlo or gradient-based optimization techniques. This method yields valid protein structures, but requires an additional P(*seq*) model, such as protein language models (Rives et al., 2021), to correct or redesign the sequence. Essentially, this approach can be viewed as optimization process of P(*structure*| *seq*) ·P(*seq*). Another approach attempts to explore the P(*backbone*) distribution. a series of protein generation models based on SE(3) invariance or equivariance networks (Jing et al., 2020; Satorras et al., 2021) have recently emerged, these method rely on an additional P(*seq*| *backbone*) process to determine the protein sequence. This optimization strategy can be regarded as P(*backbone*) · P(*seq* | *backbone*).

This step-wise design process has limitation in approximating the joint distribution through marginal distributions. The P(*structure* |*seq*) ·P(*seq*) strategy faces challenges when sampling in the high-dimensional sequence space, while the P(*backbone*) ·P(*seq* |*backbone*) strategy fails to account for explicit side-chain interactions and is bottlenecked by the capability of the fixed-backbone design model.

The ultimate goal of protein generation is to directly obtain a sequence along with its corresponding structure, i.e., to develop a model capable of describing the joint distribution P(*structure, seq*) or P(*backbone, seq*). Recently, some studies have started to adopt co-generation approaches, such as model based on co-diffusion (Campbell et al., 2024) or co-design (Ren et al., 2024). While these methods primarily rely on SE(3) networks, they still separately model the backbone and sequence, without considering side-chain conformations and leading to an insufficient description of the structure. Protpardelle (Chu et al., 2024), an all-atom protein diffusion model, similarly adopts co-generation approaches with an explicit all-atom representation, taking a step further in the field. However, the experimental results indicate that the generated sequence fails to accurately encode the intended fold, necessitating an additional round of sequence redesign and side-chain refinement.

In this study, we introduce a novel approach for all-atom protein generation called **Pallatom**. Our extensive experiments show that by learning P(*all-atom*), high-quality allatom proteins can be successfully generated, eliminating the need to learn marginal probabilities separately. To address the first critical challenge of representing side-chain coordinates for unknown amino acids in protein generation, we introduce `atom14`, a novel all-atom representation framework. This approach employs virtual atoms across all amino acid types, effectively preventing sequence information leakage while maintaining structural integrity. Inspired by AlphaFold3 (AF3) (Abramson et al., 2024), we propose a dual-track representation framework that encodes protein structures through concurrent residue-level and atom-level tokenization schemes. Central to this framework is the AtomDecoder unit, which seamlessly integrates and updates features across multiple representation spaces while simultaneously executing coordinate prediction and differentiable recycling process within a unified computational architecture. The fundamental insight underlying Pallatom is the recognition that protein all-atom coordinates intrinsically encapsulate both structural and sequential information within their spatial configuration. Directly learning P(*all-atom*) opens a new path for co-generative modeling of structure and sequence.

Our contributions are summarized as follows:

- We explore the `atom14` representation to achieve a unified description of unknown amino acid side-chain coordinates in generative tasks.
- We develop a network architecture for all-atom protein generation tasks, which effectively represents both protein backbones and sidechains.
- We use our framework to develop Pallatom, a state-of-the-art all-atom protein generative model.

## 2. Preliminaries

### 2.1. All-atom modeling and representation

Recent advances in protein structure prediction have established diverse methodologies for all-atom representation. AlphaFold2 (AF2) (Jumper et al., 2021) pioneered a frame-based approach, modeling atomic coordinates through SE(3) transformations of backbone and side-chain frames. Building upon this foundation, AF3 introduces an innovative point cloud representation that directly encodes all-atom 3D coordinates in Cartesian space. However, these representations are fundamentally inadequate for protein generation tasks due to a critical limitation: the complete absence of sequence information at the initialization stage prevents determination of the system’s total atomic cardinality. To overcome these limitations, Protpardelle introduces the `atom73` representation, which implements a quantum-inspired superposition framework. This approach stores coordinate information for all 20 possible side-chain configurations relative to aligned backbone atoms, enabling probabilistic representation of unknown amino acid types. The superposition collapses to a definitive state upon sequence determination, effectively resolving the initial uncertainty in atomic configuration.

### 2.2. Diffusion modeling on all-atom protein

The all-atom representation bypasses the complexities of SE(3) frames (Yim et al., 2023b) and Riemannian diffusion (De Bortoli et al., 2022). Gaussian-based diffusion models provide a robust theoretical framework, with EDM (Karras et al., 2022) emerging as a prominent approach. EDM has been successfully implemented in Alphafold3 and Prot-pardelle for all-atom coordinate diffusion, showcasing its efficacy in protein generation. We briefly outline EDM’s mechanism below.

Assuming the data distribution of all-atom coordinates under any representation as p_data_(**x**) with standard deviation σ_data_, the forward process involves adding Gaussian noise of varying scales to generate a series of noised distributions p_*t*_(**x**) = [*p*_*data*_ ∗ 𝒩 (**0**, *σ(t)*^2^**I**)](*x)* ≜ *p*(**x**; *σ)* with a particular configuration *σ(t)* = *t a*nd *s(t)* = 1. When 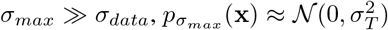 approximates pure Gaussian noise. The probability flow ordinary differential equation (ODE) is given by:

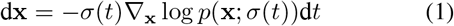

Here, ∇_**x**_ log *p(***x**; *σ)* is the score function, which does not depend on the normalization constant of the underlying density function *p*_*t*_(**x**). A neural network *D*_*θ*_(**x**; *σ)* is typically trained for each *σ u*sing the following loss function to match the score function (Song et al., 2021):

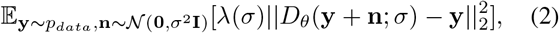

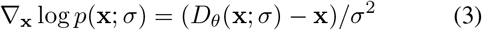

In practice, EDM introduced a preconditioning technique that stabilizes training by adjusting noise-related scaling coefficients, ensuring that the model’s inputs and outputs remain within a stable numerical range at each time step. Consequently, *D*_*θ*_ and the loss function can be derived as:

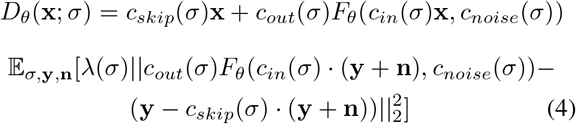

where 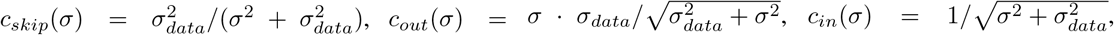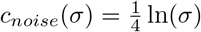 and *λ(σ)* = 1/*c*_*out*_*(σ)*^2^ represent skip scaling, output scaling, input scaling, noise conditioning, and the Coefficient respectively.

### 2.3. Framework for structure-based protein generation

Currently, the field of protein structure design lacks comprehensive methods for all-atom coordinate generation. The most relevant approaches are structure-based backbone design or sequence-backbone co-generation methods, which typically rely on a core building block to perform multi-view feature fusion and coordinate updates on protein representations (e.g., 1D single-, 2D pair-, and 3D frame-embeddings). By stacking multiple such blocks, these methods progressively refine protein structures. For instance, RFDiffusion (Watson et al., 2023), built on RoseTTAFold (Baek et al., 2021), integrates information from three tracks within each block, continuously updating and fusing representations through sophisticated network operations. Similarly, Multi-Flow (Campbell et al., 2024) and CarbonNovo (Ren et al., 2024) adopt Alpahfold2’s Invariant Point Attention (IPA) layer as their building block, augmented with additional mechanisms to refine pairwise and sequence information. Through multi-view feature fusion and coordinate updates, these methods achieve precise backbone generation.

In contrast, ProtPardelle uses a simpler U-ViT (Bao et al., 2023) architecture to generate all-atom proteins but relies on a post-processing miniMPNN module to redesign sequences and guide side-chain placement, which limits its robustness. This highlights a key limitation of simplified frameworks: their inability to effectively integrate diverse information sources. In contrast, the complex network frameworks offer significant advantages to incorporate additional auxiliary information. For example, RFDiffusion employs a self-conditioning mechanism (Chen et al., 2022) to generate preliminary structures as 2D templates for guiding sampling, while CarbonNovo leverages encoded single- and pair-features to perform sequence sampling through Markov Random Field (MRF). These strategies significantly improve the diversity and quality of generated proteins. Thus, exploring all-atom generation frameworks that integrate multiple encoding forms remains a critical research direction.

## 3. Method

### 3.1. atom14: the all-atom representation

The all-atom protein generation model faces many challenges in constructing both backbone and side-chain atoms. A pivotal initial question arises: How to represent a system with a variable number of atoms? At the initial sampling stage, both the backbone and sequence are unknown, however, the atom number of a system depends on unique sequence, once the sequence is determined, it also dictates the structure.

To avoid potential conflicts arising from the simultaneous design of sequence and structure, we introduce the `atom14` representation, establishing a unified framework through virtual atom integration that normalizes heavy atom counts across all amino acid types. This padding strategy that empirically positions additional virtual atoms to align with the C*α* coordinates of each amino acid residue. Through this representation, a protein with *N r*esidues, denoted as 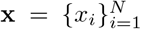, is transformed into a fixed-size point cloud **x**^0^ ∈ℝ^*N ×*14*×*3^. For example, if residue *x*_*i*_ is `CYS`, its coordinates [N, C_*α*_, C, O, C_*β*_, S_*γ*_] ∈ℝ^6*×*3^ are augmented with eight virtual atoms aligned at the C_*α*_ position, yielding 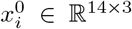. We denote the system as 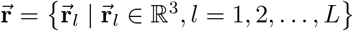 with *L* = *N* × 14 and define the noise level as 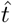.

The primary task of all-atom generation involves learning the score function 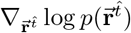 through a diffusion process that progressively perturbs the initial state 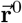 at each noise level 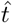. While the virtual atom framework inherently precludes explicit element-specific assignments, we posit that the all-atom coordinate distribution sufficiently encodes essential sidechain properties, including hydrophobicity, polarity, hydrogen bonding, and salt bridge formation. Therefore, we introduce an auxiliary visualization module that predicts amino acid types. With discarding the redundant virtual atoms based on the predicted amino acid type as a post-processing step, we can generate all-atom proteins with corresponding sequences from a 3D point cloud noise. Details are presented in Figure 3 and Appendix C.1

### 3.2. MainTrunk: the denoising network

We refer to the protein generation task as the generation of all-atom coordinates. In the `atom14` representation, a protein with *L a*toms can be expressed as 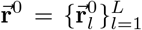, which 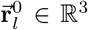 represents an atom coordinate. For each noise level 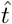 in the diffusion process, the network predicts the updated coordinates 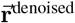 from the input 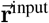.

The network comprises two main components: a feature encoder and iterative decoding units. Figure 1A illustrates the main architecture. We implement a dual-track representation framework, employing local attention mechanisms for atomic-level feature extraction and global attention mechanisms for residue-level feature integration.

**Figure 1.**
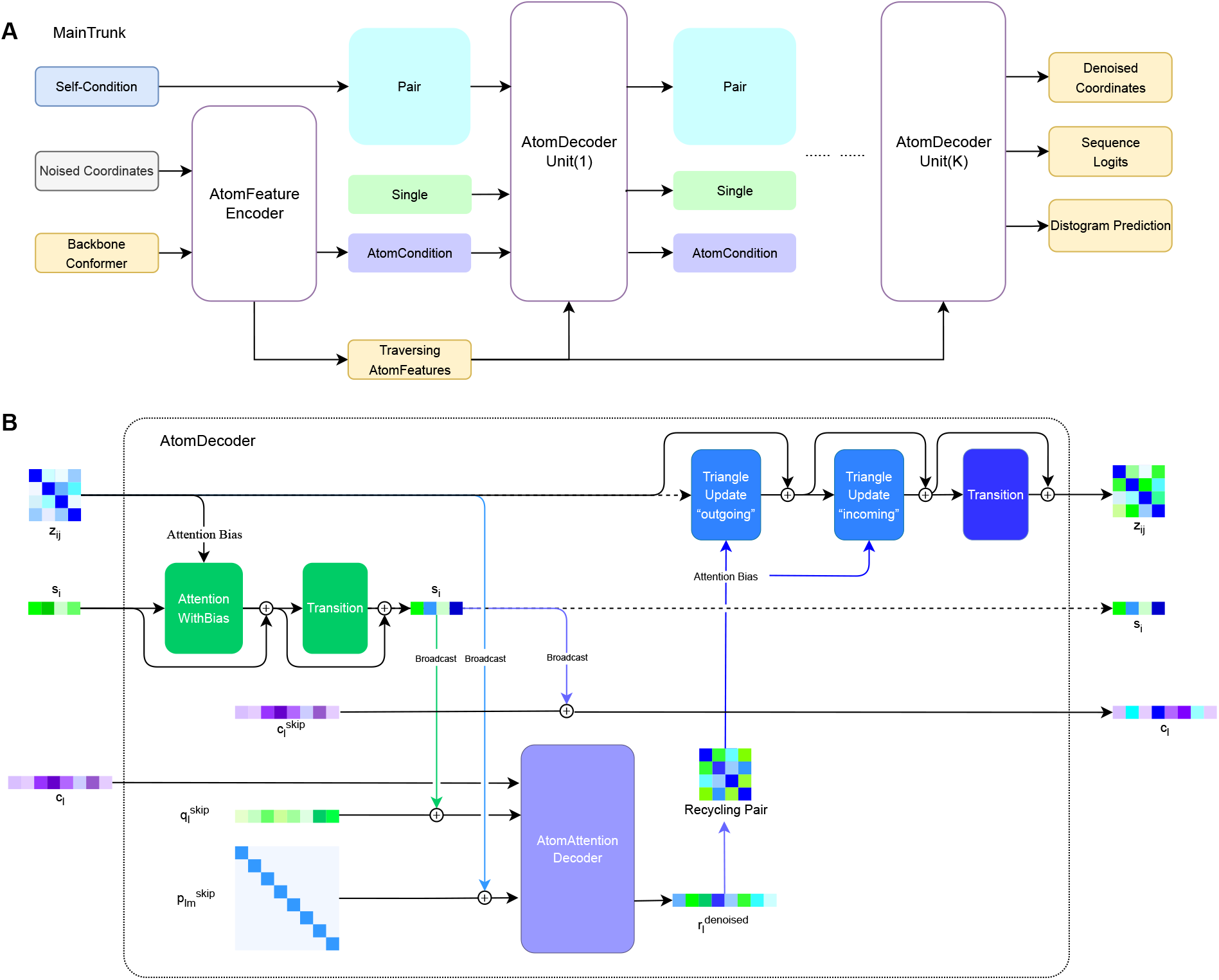
Pallatom model framework: (A, B) Architecture of the network and the AtomDecoder unit.

For feature initialization and encoding (Algorithm 4), We first initialize “traversing” atomic embeddings (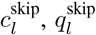and 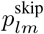 by integrating three key components: (1) structural features from alanine’s standard conformation to establish a stable backbone frame, (2) residue positional encoding, and (3) noisy coordinate vectors. For residue-level embedding, single embedding (*s*_*i*_) is initialized using diffusion timestep encoding and residue positional encoding, while pair features (*z*_*ij*_) incorporate relative positional encoding and self-condition template distograms from previous predictions. Detailed features are recorded in Appendix Table 5.

In the decoding phase, we implement an iterative refinement mechanism that facilitates bidirectional information flow: residue-level embedding are propagated to atomic-level, while atomic information is subsequently integrated back through a differentiable recycling process. These operations are encapsulated within an AtomDecoder unit (Figure 1B), which is designed to continuously update the coordinates. In practical implementation, we identify a fundamental challenge in the decoding architecture: preserving residual connections between residue-level and atomic-level embeddings across successive decoder units may result in significant information redundancy, especially during the residue-level feature broadcasting process (where “feature broadcasting” refers to the process similar to AF3’s approach of replicating residue-level features and incorporating them into corresponding atom-level features). A comprehensive analysis of this phenomenon and our solution is provided in Appendix C.5. To address this challenge, the three “traversing” atomic embeddings serve as stable information carriers, facilitating efficient residue-level propagation while enabling coordinate refinement through the accumulated 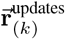 in the *k-*th unit.

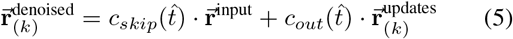

Following coordinate denoising, the partially denoised positions 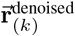 are transformed into relative distance matrices and subsequently recycled into the *z*_*ij*_ *p*air embeddings. Concurrently, the triangle update module selectively maintains physically valid triangular constraints while eliminating geometrically inconsistent features, enabling iterative improvement of structural quality throughout the decoding process. Details can be found at Appendix Algorithm 2.

### 3.3. SeqHead: sequence decoder

The `atom14` representation, while effective for structural denoising, inherently lacks elemental information in the generated side chains, preventing direct extraction of protein sequence information from the atomic coordinates alone. To bridge the gap between structural generation and biological interpretation, we introduce a `SeqHead` module that converts the predicted atomic coordinates into corresponding amino acid sequences, enabling the transformation of geometric data into biologically meaningful protein structures. Specifically, we aggregate the atomic embedding (*q*_*l*_) corresponding to each residue and process them through a linear transformation layer to predict probability distributions over the 20 canonical amino acid types **â** _(*k)*_ ∈ℝ^*N ×*20^. We take the predictions from the last unit as the final sequence logits, **â** = **â**_(*K)*_.

### 3.4. Training Loss

Our training method mainly follows the application and improvements of the EDM framework. The denoising all-atom positions score-matching losses are described according to Eq.(2). Given that the network architecture lacks inherent equivariance constraints and iteratively refines coordinates across multiple decoding stages, it is imperative that the coordinates transformations between these stages remain invariant under changes in orientation to ensure consistent geometric interpretations. Therefore, we employ an aligned MSE loss. We first perform rigid alignment of the ground truth 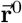 on the denoised structure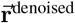 using the Kabsch algorithm, yielding 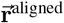. The MSE loss is then defined as: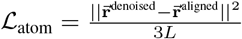

For sequence decoding, we use the standard cross-entropy loss function to evaluate the difference between the predicted sequence **â** and the true sequence **a**_0_. The loss function is defined as ℒ_seq_ = CE(**â, a**^0^). The primary loss function of the network is:

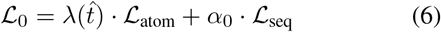

To capture the fine-grained characteristics of the allatom structure, we introduce the simplified smooth local distance difference test (LDDT) loss from AF3 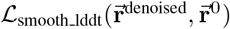. This can be found in the Appendix algorithm 8. Additionally, we implement two distogram-based loss function ℒ_dist___res_ and ℒ_dist___atom_ to enforce global/local distance constraints at the residue/atomic level, ensuring proper maintenance of overall protein topology throughout the optimization process. We introduce an intermediate layer loss ℒ_*med*_, that supervises the sequence and structure predictions from each AtomDecoder unit, significantly enhancing both model performance and inference stability through progressive refinement. Details of training losses can be found in Appendix D. The total loss can be written as:

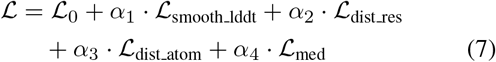

#### Algorithm 1

Pallatom Inference

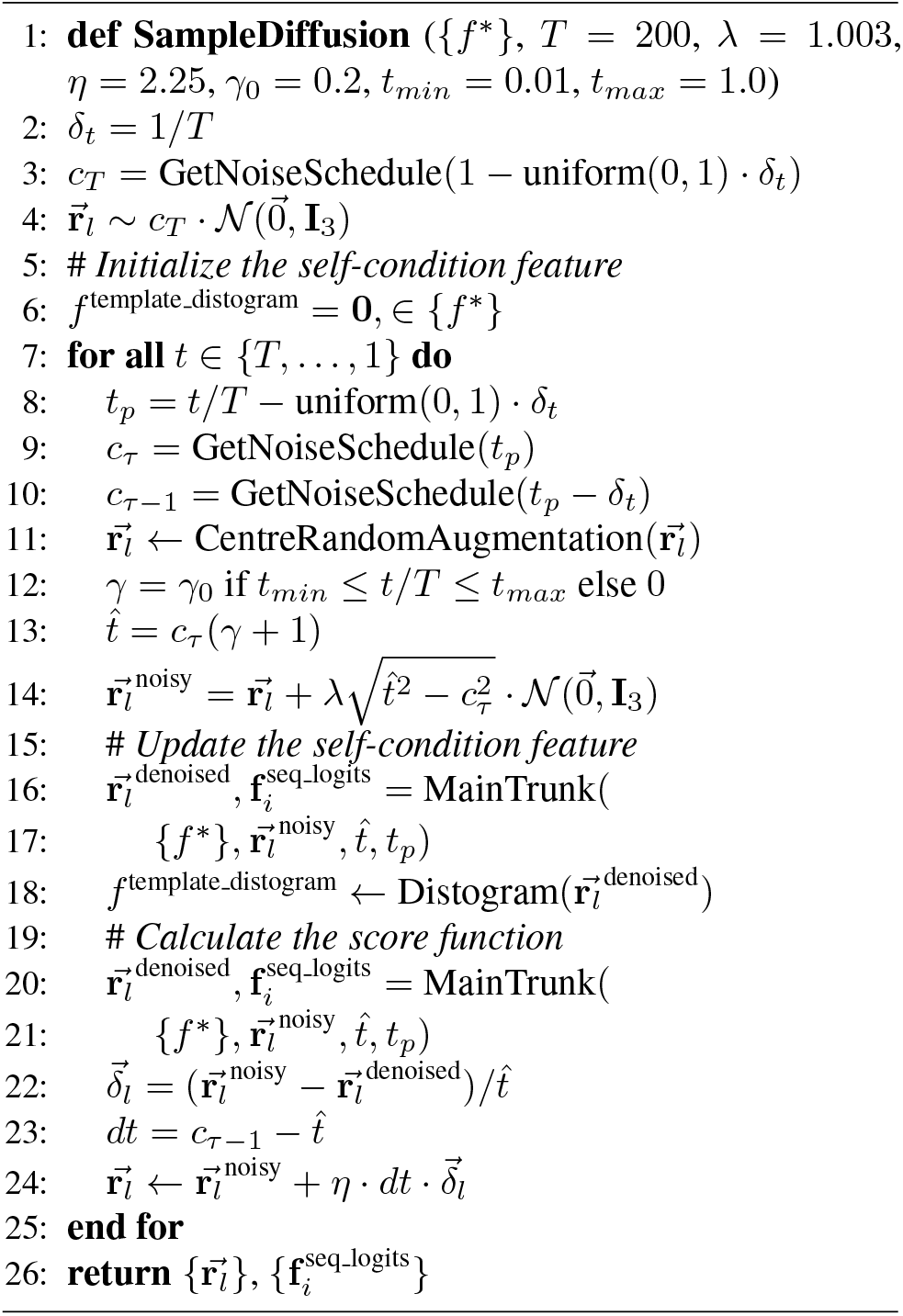

### 3.5. Sampling

The sampling process is described in Algorithm 1. The initial atoms are sampled from a Gaussian distribution on ℝ^*L×*3^. We only use the first-order Euler method as the ODE solver with *T* steps for discretization. Optionally, additional noise can be injected during the sampling steps to introduce stochasticity into the ODE solving process. We focus only on the sequence distribution decoded by the network in the final sampling step, employing a low-temperature softmax strategy to derive an approximate discrete one-hot amino acid sequence as the final sequence.

## 4. Experiments

### 4.1. Training setting

The training dataset of the model includes the PDB (Zardecki et al., 2022) and AlphaFold Database (AFDB) (Varadi et al., 2021). We performed rigorous data cleaning on augmented data from AFDB to obtain high-quality results. Details can be found in the Appendix B. We focus on small monomer proteins that can be easily synthesized using commercial oligo-pool method and the models are trained on crops of lengths up to 128. The model training utilized the Adam optimizer (Kingma & Ba, 2017) with a learning rate of 1e-3, *β*_1_ = 0.9, *β*_2_ = 0.999, and a batch size of 32. Details are provided in the Appendix Table 6.

### 4.2. Metrics

While some evaluation criteria used for protein backbone generation are not suited for the new task, we propose new metrics specifically designed for assessing all-atom protein generation.

The first criterion is structure designability. The self-consistency process assesses the designability of protein backbones (**DES-bb**). This involves using a fixed-backbone design model (e.g., ProteinMPNN (Dauparas et al., 2022)) to generate *N*_*seq*_ *s*equences for the backbone, which are then folded by structure prediction models like ESMfold (Lin et al., 2023). The backbone’s designability is evaluated by the optimal TM-score or C_*α*_-RMSD between the folded and original backbones. However, this metric is not suitable for all-atom proteins, which include side-chain atoms. Therefore, we similarly define the designability of all-atom protein generation, denoted as **DES-aa**. For the all-atom proteins, the sequence is used to predict the structure, and the sample is considered designable if the mean pLDDT of the predicted structure exceeds 80 and the all-atom RMSD (aaRMSD) is less than 2 *Å*. This metric ensures atomic-level accuracy and provides strong confidence in the structural integrity and designability of the predicted protein.

The second criterion is structure diversity, denoted as **DIV-str**. This can be quantified by calculating the clusters number of the designable structures using Foldseek (Van Kempen et al., 2024). For all-atom proteins, we use a similar diversity evaluation method for the generated sequences, denoted as **DIV-seq**. Specifically, we use MMseq2 (Steinegger & Söding, 2017) to calculate the clusters number of the designable sequences.

The last criterion is structure novelty, which evaluates the structural similarity between the generated backbones and natural proteins in the PDB, denoted as **NOV-str**. This is calculated by the TM-score of the generated designable backbones compared to the most similar proteins in the PDB. In our evaluation, we use the following two modes:

- CO-DESIGN 1: For methods that can predict both all-atom coordinates and sequences, **DES-aa** is used. Diversity and novelty are evaluated based on the allatom designable proteins.
- PMPNN 1: Other methods use **DES-bb** with *N*_*seq*_ *=* 1. Specifically, we calculate and display the two **DES-bb (w/wo)** under conditions with and without the pLDDT > 80 constraint. Diversity and novelty are evaluated based on the proteins filtered by **DES-bb (w)**.

### 4.3. Results

We sample Pallatom with 200 time steps using a noise scale *γ*_0_ = 0.2, a step scale *η* = 2.25 and evaluate 250 proteins sampled for each length *L =* 60, 70, 80, 90, 100, 110, 120. Our primary comparisons are with state-of-the-art methods capable of generating all-atom proteins, such as Protpardelle and ProteinGenerator (Lisanza et al., 2023). For backbone generation, we compare with RFdiffusion. We also compared Multiflow, which is capable of generating both backbones and sequences. All methods are evaluated using their open-source code and default parameters.

As shown in Table 1 and Figure 2A, Pallatom demonstrates superior performance in the CO-DESIGN 1 benchmark for all-atom protein generation. Remarkably, despite lacking specific training on fixed-backbone design tasks, Pallatom achieves comparable results to ProteinMPNN by predicting sequences with a single linear layer derived from aggregated atomic embedding. Comprehensive analysis of sequence quality between our design and ProteinMPNN, including detailed performance metrics, is presented in the Appendix G.4. These results substantiate Pallatom’s capacity for all-atom structure design, validating our hypothesis that modeling *P(all-atom*) effectively captures the fundamental relationship between protein structure and sequence. Notably, Pallatom generates all-atom protein structures with substantially enhanced structural diversity, outperforming existing methods by achieving significantly higher diversity than both Multiflow and ProteinGenerator, while simultaneously attaining the highest sequence diversity across all evaluated approaches.

**Table 1.** Comparison of various methods. Protpardelle utilized ProteinMPNN as an auxiliary tool in all-atom proteins generation, resulting in identical results for CO-DESIGN 1 and PMPNN 1. In the case of Multiflow, which can only generate protein backbones and sequences without side chains, the reported DES-aa metric is based on C_*α*_RMSD rather than aaRMSD.

| Method | CO-DESIGN 1 |  |  | PMPNN 1 |  |  |
| --- | --- | --- | --- | --- | --- | --- |
| | DES-aa (↑) | DIV-str/seq (↑) | NOV-str (↓) | DES-bb (w/wo) (↑) | DIV-str (↑) | NOV-str (↓) |
| Protpardelle* | 30.00% | 15 / 26 | 0.747 | 30.00% (80.23%) | 15 | 0.747 |
| ProteinGenerator | 43.14% | 83 / 706 | 0.791 | **93.14% (96.34%)** | 151 | 0.785 |
| Multiflow* | 62.74% | 134 / 1042 | 0.753 | 84.69% (95.43%) | 184 | 0.744 |
| RFdiffusion | N/A | N/A | N/A | 78.29% (84.40%) | 165 | 0.805 |
| Pallatom | **85.03%** | **291 / 1466** | **0.719** | 89.89% (94.46%) | **318** | **0.716** |

**Figure 2.**
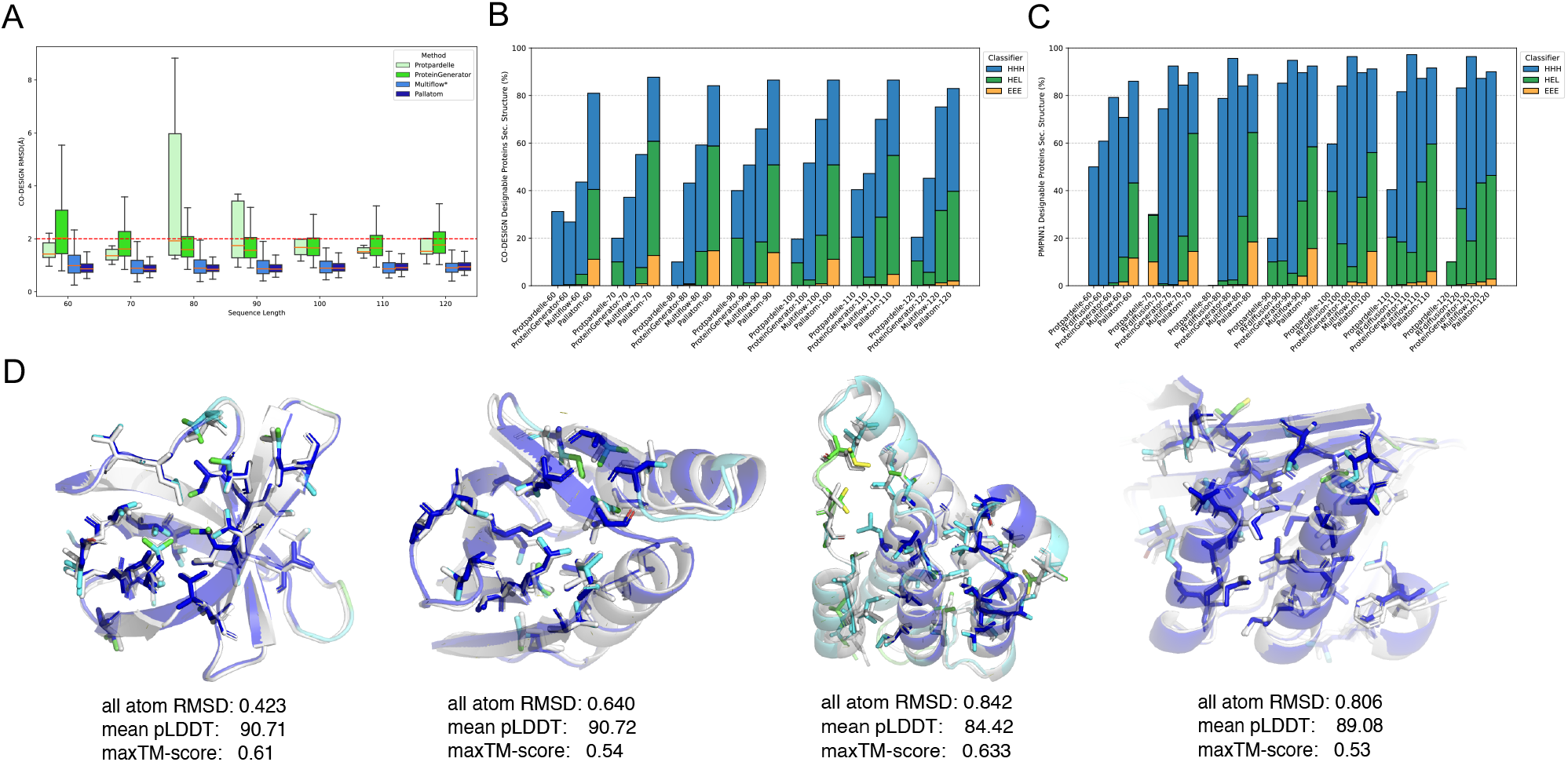
Evaluation of proteins sampled from Pallatom. (A) Boxplot of aaRMSD for proteins sampled by various methods under the CO-DESIGN 1 mode. Multiflow exhibits the C_*α*_RMSD. (B, C) The proportions of secondary structures in designable proteins across different lengths are presented for CO-DESIGN 1 and PMPNN 1 modes across various methods, with the total height of the y-axis representing the designability. (D) Examples of high-quality, novel all-atom proteins sampled by Pallatom.

**Figure 3.**
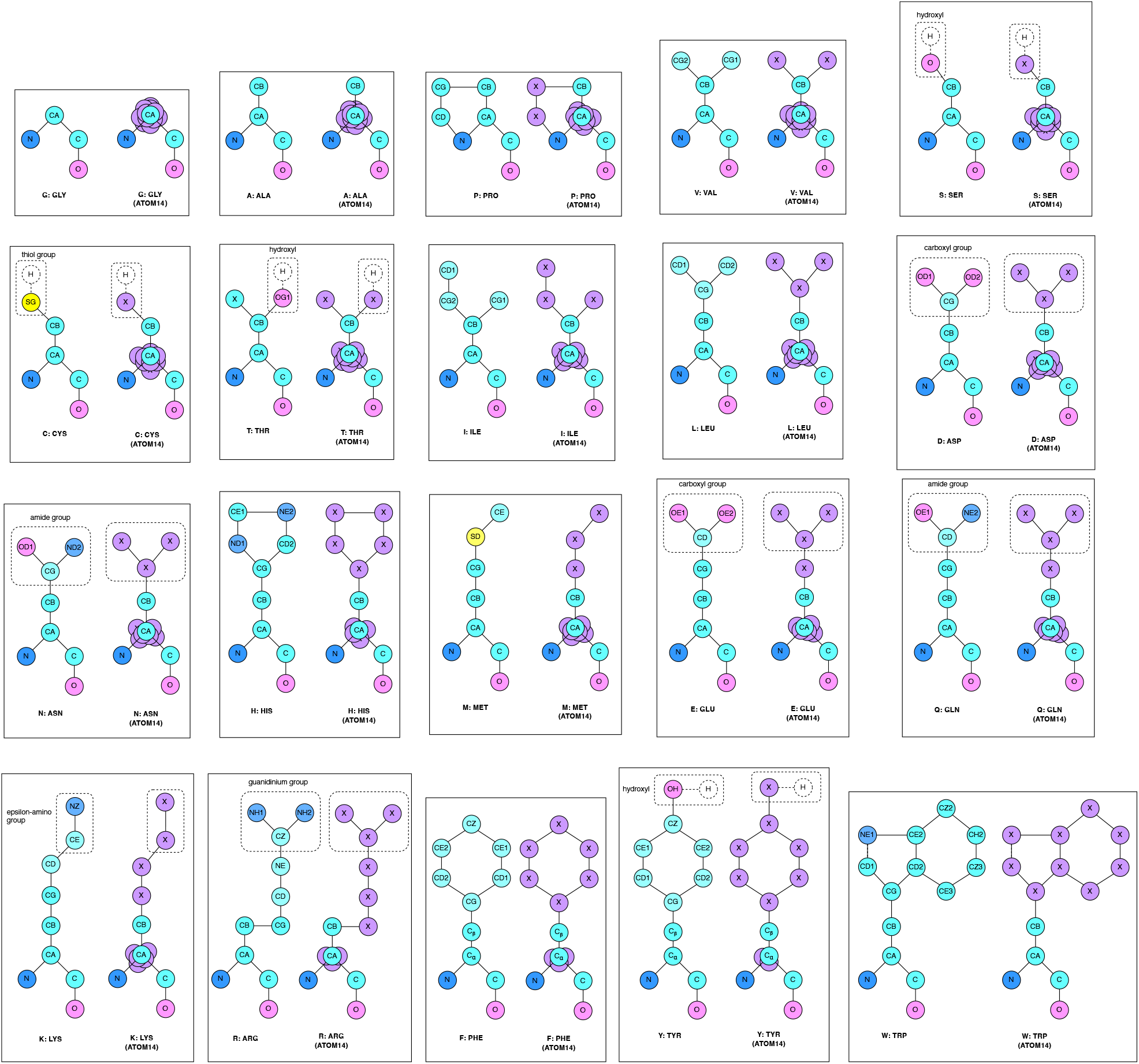
`atom14` encoding schemes.

In the PMPNN 1 benchmark, we observe that the DES-bb(wo) metric systematically overestimates designability relative to the pLDDT-constrained DES-bb(w) metric, as demonstrated by substantial discrepancies in Protpardelle’s designability evaluations, highlighting the importance of quality constraints in designability assessment. The backbone generative models like ProteinGenerator, Multiflow and RFdiffusion demonstrate comparable performance. However, Protpardelle demonstrates suboptimal performance across all evaluation metrics, underscoring the critical importance of both network architecture design and atomic representation strategies in developing effective generative models for protein design. Consistent with its CO-DESIGN 1 performance, Pallatom demonstrates robust designability while simultaneously generating structurally diverse and novel protein conformations, effectively balancing these traditionally competing objectives. A detailed analysis in Figure 2BC shows Pallatom’s stable performance across protein lengths and balanced secondary structure distributions, in contrast to the preference of comparative methods for specific secondary structures like *α*-helical ones. The case studies in Figure 2D highlight Pallatom’s ability to design highly ordered proteins with hydrophobic cores and hydrophilic surfaces, consistent with high pLDDT structures predicted by ESMFold, underscoring its understanding of protein folding principles.

To assess generalization capabilities, we extensively evaluated Pallatom’s performance on out-of-distribution (OOD) protein lengths, conducting systematic sampling and analysis across a broad range of 150-400 residues. The results demonstrate that Pallatom exhibits exceptional scalability, achieving the highest designability even at over twice the maximum training length (*L* = 128). We also analyze secondary structure preferences in generated proteins across different methods, particularly for longer sequences, with comprehensive distribution statistics and detailed comparisons provided in the Appendix G.1. We also performed a statistical analysis of sampling duration, which demonstrates the efficiency of Pallatom. The results are presented in Appendix G.7.

### 4.4. Hyperparameter

We analyzed the impact of the sampling parameters and Table 2 presents the metrics for Pallatom when sampling 250 proteins with *L* = 100. We observed that, under the same noise scale, increasing the step scale *η* leads to a corresponding rise in designability, as well as an increase in the proportion of all-helix structures within the secondary structure. However, this improvement in designability comes at the cost of reduced structural diversity, indicating a trade-off between these two metrics. We further demonstrated the impact of varying step scales on the sampling of OOD-length proteins, with detailed results provided in the Appendix G.3. Furthermore, we found that reducing the additional noise level to *γ* = 0.1 slightly enhances designability but also significantly decreases structural diversity, consistent with the findings of previous work (Yim et al., 2023b).

**Table 2.** Pallatom sample metrics.

|  |  |  |  |  |  |
| --- | --- | --- | --- | --- | --- |
| **Noise Level** $\gamma_0$ | 0.2 | 0.2 | 0.1 | 0.2 | 0.2 |
| **Step Scale** $\eta$ | 1.75 | 2.25 | 2.25 | 2.75 | 3.25 |
| $N_{\text{steps}}$ | 200 | 200 | 200 | 200 | 200 |
| **DES-aa** (↑) | 57% | 87% | 89% | **94%** | 93% |
| **DIV-str** (↑) | 56 | **64** | 52 | 55 | 46 |
| HHH(%) | 28% | 28% | 45% | 35% | 38% |
| HEL(%) | 56% | 54% | 44% | 47% | 51% |
| EEE(%) | 16% | 18% | 11% | 18% | 11% |

### 4.5. Ablation Studies

Building upon the hypothesis that all-atom coordinates intrinsically encode essential protein information, we systematically investigated the key factors underlying Pallatom’s superior designability performance. Table 3 presents the results, with the first row (atom14) serving as the baseline. We first conducted an ablation study on the `atom14` representation. For comparative analysis, we implemented the `hybrid14` representation and retrained the model. Similar to Protpardelle’s methodology, the `hybrid14` representation maintains a superposition of 20 possible side-chain configurations, which collapse into a single hybridized residue state based on predicted sequence probabilities during the generation process. However, as shown in the second row, the sequence-guided all-atom approach not only failed to improve performance but actually resulted in reduced sequence quality (lower DES-aa) and compromised backbone generation (decreased DES-bb), revealing fundamental limitations in this design strategy. We hypothesize that this phenomenon stems from the absence of a well-defined mapping between noisy coordinate space and sequence space during diffusion. The sequence predictions derived from noisy structural intermediates exhibit substantial deviations from native foldable sequences, and these discrepancies are amplified throughout the reverse diffusion process, ultimately hindering the generation of biologically plausible sequences.

**Table 3.**
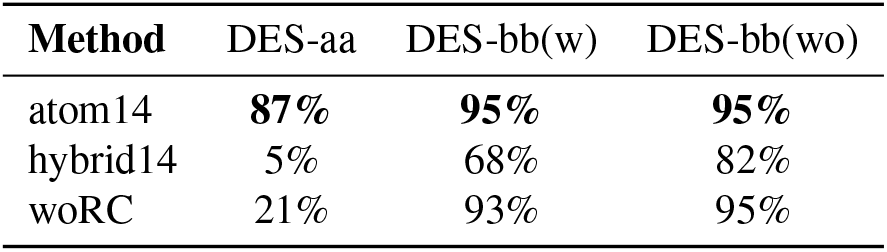
Ablation Studies evaluating the key concept of Pallatom to designability.

| **Method** | DES-aa | DES-bb(w) | DES-bb(wo) |
| --- | --- | --- | --- |
| atom14 | **87%** | **95%** | **95%** |
| hybrid14 | 5% | 68% | 82% |
| woRC | 21% | 93% | 95% |

We next assessed the impact of the differentiable recycling process (woRC, shown in the third row). Our evaluation indicated that while the recycling process had little effect on backbone quality (DES-bb remained almost unchanged), it significantly impacted sequence generation, leading to a substantial reduction in DES-aa. These results highlight the crucial role of the recycling process in the components and its differential influence on various aspects of protein design. The detailed setup of the ablation studies can be found in the Appendix E.

## 5. Discussion

We introduce Pallatom, a highly efficient end-to-end all-atom protein generation framework that simultaneously captures the relationship between sequence and structure, enabling state-of-the-art performance. Our proposed all-atom protein `atom14` representation eliminates the constraints of explicitly defining all amino acid types during generation, offering a more precise approach for representing coordinates in all-atom systems. We have redesigned a fundamental unit capable of fusing, recycling and updating diverse features of all-atom proteins while predicting their coordinates. The new framework efficiently adapts to all-atom protein structure diffusion generation. The results demonstrate that models learning P(*all-atom*) exhibit strong performance and diversity in *de novo* protein generation, unlocking new pathways for protein design. Future work includes developing a more generalized model architecture and extending our framework to enable the design of complex systems.

## Impact Statement

The novel protein generation method proposed in this paper, which is based on learning the distribution of all-atom coordinates, aims to advance the paradigm of protein design engineering. This work holds significant potential for broad societal impact and direct applications. For instance, it offers greater utility in tasks such as enzyme design, drug design involving small molecule binding pockets, and other applications requiring all-atom protein constraints.

## Code Availibility

The Pallatom is available on GitHub (https://github.com/levinthal/Pallatom).

## A. Related work

### Backbone Generation

Diffusion models based on SE(3)-equivariant network architectures and protein representations using rigid frames have achieved significant success in protein generation, as evidenced by models like Chroma (Ingraham et al., 2023), Genie2 (Lin et al., 2024), RFdiffusion (Watson et al., 2023), FrameDiff (Yim et al., 2023b), FrameFlow (Yim et al., 2023a), Proteus (Wang et al., 2024), and FoldFlow2 (Huguet et al., 2024). These models now support multi-condition controlled generation and have been extensively validated through both in-silico and wet-lab experiments.

### Codesign Models

Recent methods have explored a co-design approach that simultaneously designs both the backbone and sequence. Multiflow (Campbell et al., 2024) utilizes a diffusion process that jointly operates on discrete sequences and continuous SE(3) backbones, eliminating the need for sequence redesign with ProteinMPNN (Dauparas et al., 2022). CarbonNovo (Ren et al., 2024) employs a similar approach, using SE(3) diffusion on the protein backbone while simultaneously designing a sequence at each step with the MRF decoder. The sequence information is then embeded using the protein language model ESM2-3B (Rives et al., 2021) to guide structural generation.

### All-Atom Generation

Recent research teams have begun exploring fully all-atom generative representation systems. For instance, Protpardelle (Chu et al., 2024) employs a coordinates diffusion model to support the generation of all-atom protein structures. ProteinGenerator (Lisanza et al., 2023) applies Euclidean diffusion on one-hot encoded sequences, combined with a structure prediction module to obtain all-atom structure. Similarly, PLAID (Lu et al., 2025) achieves sequence design through latent space diffusion within ESMFold while decoding full-atom protein configurations. RFdiffusionAA (Krishna et al., 2024), based on fine-tuning the RoseTTAFold2 (Baek et al., 2023), can produce backbone structures of proteins and small molecule complexes but lacks side-chain conformations for standard amino acids. We focus on the methods directly generate all-atom protein structures, the most relevant work is Protpardelle, which uses an end-to-end approach to generate all-atom structures.

Beyond protein modeling, recent works leverage atomic-level representations for localized design tasks. For instance, Abdiffuser (Martinkus et al., 2023) introduces a universal four-atom side-chain template for antibody CDR redesign, preserving dihedral freedom via pseudo-carbon atoms while integrating ideal amino acid templates for rotamer construction. FAIR (Zhang et al., 2023) adopts a two-stage approach for protein pocket design: initial backbone/sequence generation followed by iterative refinement to ensure sequence-side-chain consistency. PocketFlow (Zhang et al., 2023) extends this by simultaneously designing pocket sequences and all-atom structures through flow matching across backbone, side-chain torsion angles, and sequences. PepFlow (Li et al., 2024) similarly utilizes multi-modal flow matching (torsion angles + sequences) for peptide design. However, such decoupled representations risk sequence-structure conflicts and steric clashes. Pallatom addresses these limitations through its `atom14` representation, which intrinsically fuses structural and sequential modalities to minimize explicit conflicts.

## B. Training Datasets

### B.1. PDB Data

We used PISCES (Wang & Dunbrack Jr, 2003) to obtain the required PDB list. For training, we selected a subset of PDB entries with a resolution of <3Å and a 95% sequence identity threshold. We then performed standard filtering to remove any proteins with >50% loops and applied a series of folding quality filters (described below). This process resulted in 7,459 structures.

### B.2. Augmented Data

Augmented data are widely used in protein modeling-related work. Consequently, we supplemented our dataset with the AlphaFold Database (AFDB) (Varadi et al., 2021). The AF2 predicted structure database is available under a CC-BY-4.0 license for both academic and commercial uses. The AFDB contains 214 million data points, while (Barrio-Hernandez et al., 2023) provides a redundancy-reduced version (AFDB-cluster) through sequence-structure clustering. We performed additional curation on AFDB-cluster, applying an average pLDDT threshold ( ≥80) and maximum sequence length restriction (128 residues) yielded 582,652 refined structures.

Furthermore, we implemented refined filtering strategies to acquire designable and high-quality data by evaluating two structural metrics — Average Neighboring Density and Core Residue Ratio — for retaining proteins with superior packing quality. The curation workflow operates as follows:

Average Neighboring Density: Specifically, we used a 10 Å distance cutoff for neighbor identification. The neighbor count is averaged across the entire protein length. This serves as a packing quality filter. The threshold values were empirically determined—we adopted 20.0 as the cutoff for Pallatom training. Note that this threshold may require further adjustment for protein datasets of varying lengths.

Core Residue Ratio: We calculated the solvent-accessible surface area (SASA) for each residue using DSSP (Touw et al., 2015), with residues having SASA ≤ 0.2 classified as core residues. A final cutoff of ≥30% core residue content was applied to retain high-quality training data.

We found that these two filters effectively selected training data with satisfactory packing quality.

Furthermore, the number and distribution of secondary structures can define the compactness and diversity of folding. We used DSSP (Touw et al., 2015) to assign the protein secondary structures and removing structures with loop content >50%. We also apply filter on the total number of continuous segments of *β*-sheet and *α*-helix to maintain structural diversity. Specifically, we define continuous segments of *β*-sheet and *α*-helix, where a continuous segment must contain at least 2-3 consecutive residues of the same secondary structure. The total number of continuous segments in the entire protein is then calculated. For Pallatom training, we applied a cutoff of 4 continuous segments as a filtering threshold during data processing. This step helps further remove overly simplistic secondary structures from the AFDB-cluster dataset.

For highly extended structures, we limited the radius of gyration (*R*_*g*_) to less than 25.0. To avoid overly long continuous unstructured regions within the protein structures, we restricted the maximum length of each loop to 15. Finally, we used the FoldSeek easycluster algorithm to remove redundant structures, setting a TM score threshold of 0.8 and a coverage of 0.9, which removed approximately 30% of highly similar structures. After applying these stringent filters, only 27,697 protein structures remained. These structures typically exhibit good folding and high designability.

## C. Algorithms

We present a comprehensive description of Pallatom’s all-atom representation `atom14`, architectural modules and algorithmic workflow. In the accompanying pseudocode, components highlighted in blue maintain substantial similarity to AlphaFold3’s implementation and are not expanded to prevent redundancy while ensuring clarity.

### C.1. atom14

To resolve the discrepancy in atom counts across amino acids within an all-atom diffusion model, we designed an atomistic representation called `atom14`. As shown in Figure 3, this representation differs from conventional amino acid encoding schemes. In the `atom14` framework, the side chains of all amino acids are uniformly represented using 14 atom coordinates.

For atoms absent in the native conformation of a given amino acid, their coordinates are overlapped with a reference atom position, specifically the C*α* atom in the Pallatom implementation. For example, glycine (Gly), which only contains four backbone atoms (C, C*α*, N, and O), has the remaining 10 “excess” side-chain positions all assigned to the C*α* coordinate. To avoid pre-specifying atom types, the `atom14` representation retains elemental types only for backbone atoms. All side-chain atoms are uniformly predefined as masked types, regardless of their actual chemical identities. It is important to note that this unified representation introduces degeneracy: certain amino acids share similar side-chain configurations (e.g., Asn vs Asp, Glu vs Gln, Cys vs Ser, Thr vs Val). Consequently, the final model output must incorporate an additional mechanism to resolve ambiguities in amino acid type determination arising from this degeneracy.

### C.2. MainTrunk

Algorithm 2 details the denoising process of the Pallatom `MainTrunk` network.

### C.3. TemplateEmbedder

In the TemplateEmbedder module, we retained only the necessary mask and distogram features, and additionally included the timestep feature for self-conditioning.

#### Algorithm 2

MainTrunk

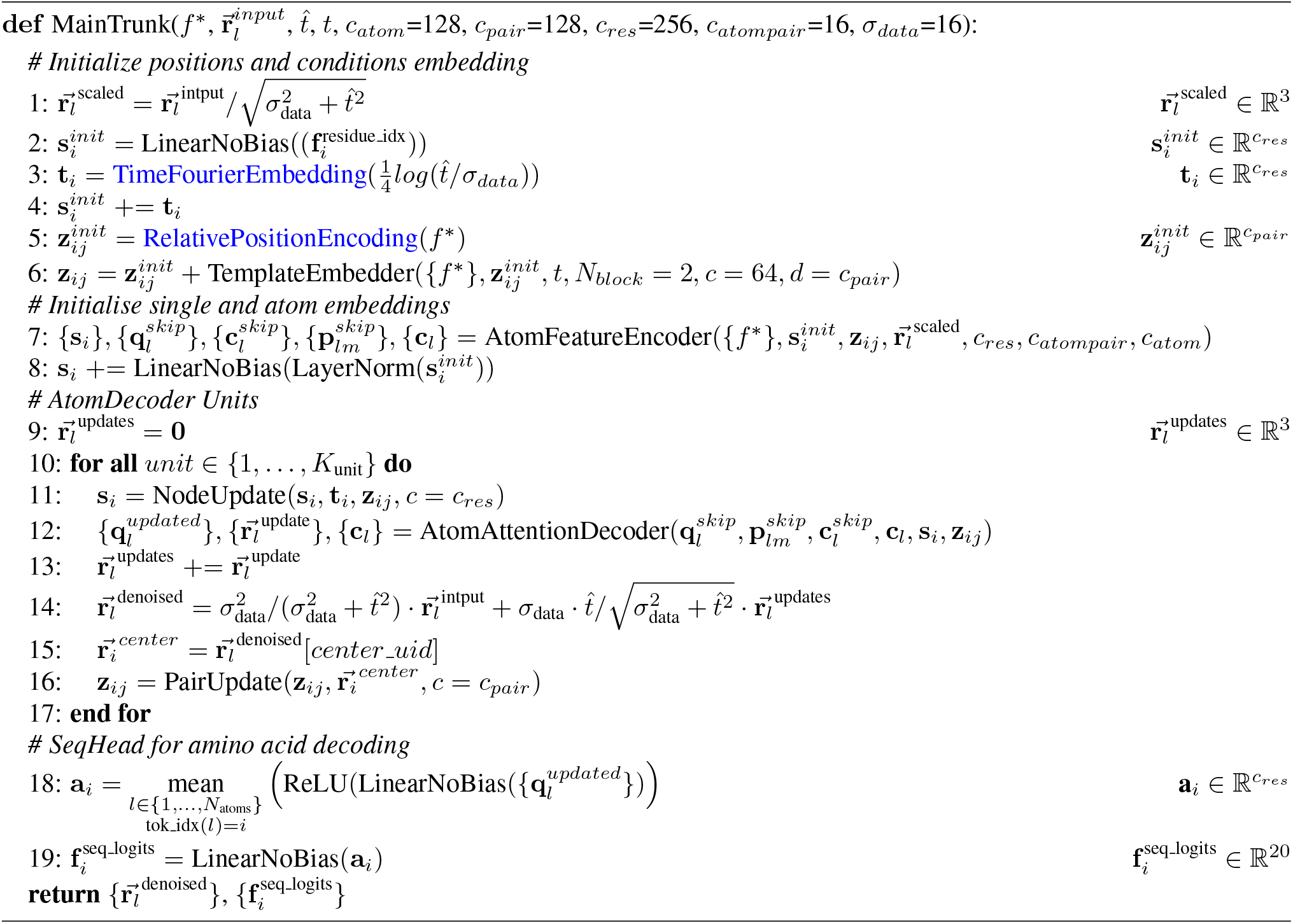

#### Algorithm 3

Template Embedder

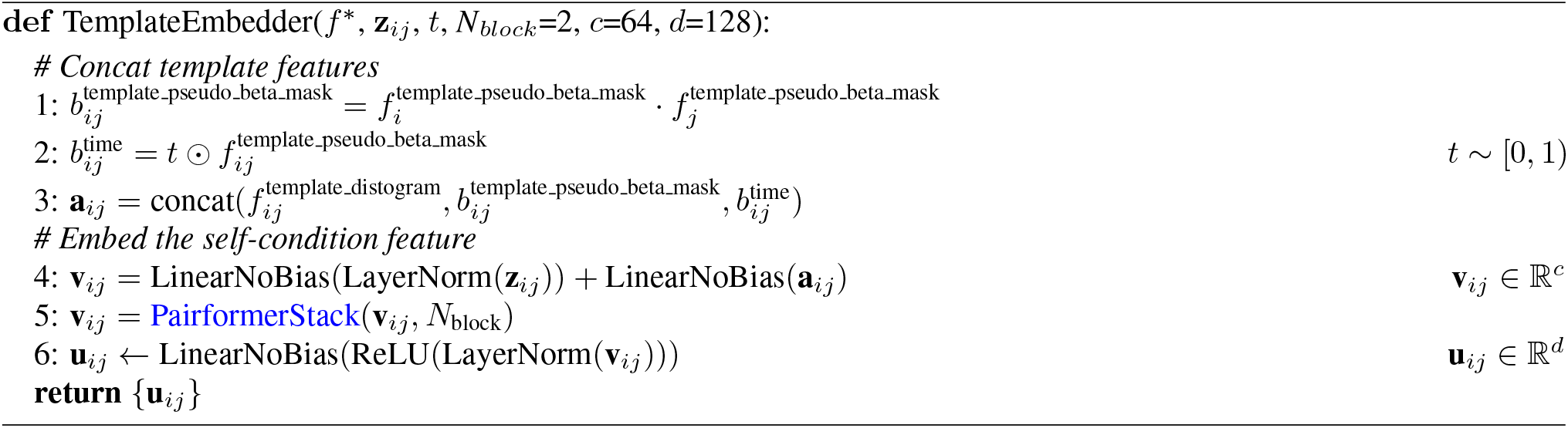

### C.4. AtomFeatureEncoder

The AtomFeatureEncoder is adapted from AF3, with modifications for all-atom generation. We exclusively utilize alanine as the reference conformer, eliminating the need for predefined amino acid sequence information during the initialization phase. Furthermore, we standardized residue representations using the `atom14` format, with virtual atoms to prevent information leakage. The initialization process for traversing atomic embeddings is detailed as follows:

#### Algorithm 4

AtomFeatureEncoder

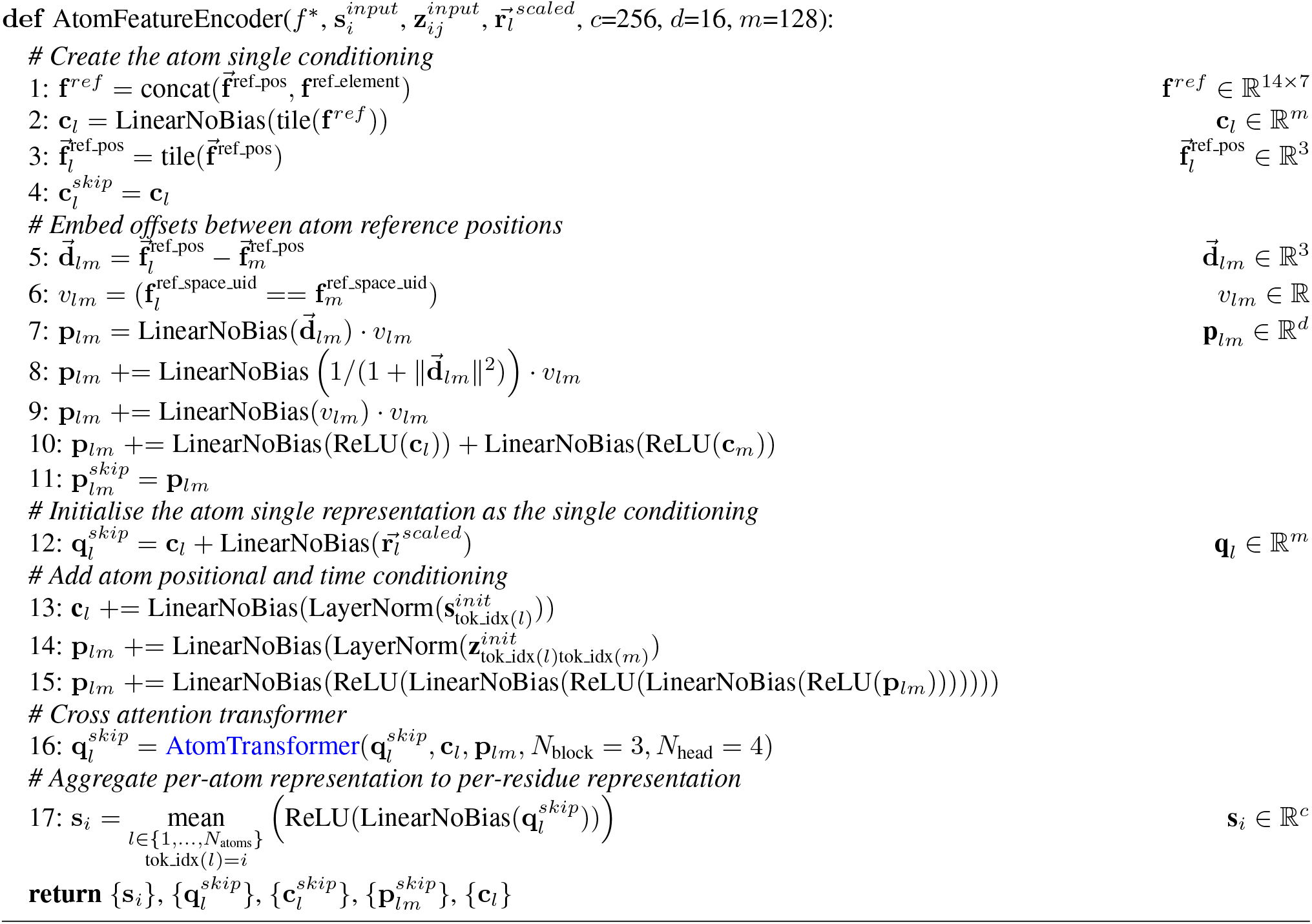

### C.5. AtomAttentionDecoder

We developed a novel information management strategy for decoder layers that simultaneously integrates updated single- and pair-embeddings into traversing atomic-level embeddings. This approach effectively prevents redundant structural information accumulation while maintaining essential feature propagation.

We present a mathematical derivation to analyze information redundancy arising from residual connections between residue-level and atomic-level embeddings, identifying key factors contributing to “traversing” strategies. For notational clarity, we define *s*_*i*_ *a*s the residue-level embedding and *q*_*i*_ *a*s the atom-level embedding for the *i-*th unit, with broadcast(*s*_*i*_) denotes the operation that propagates residue-level information to atomic-level. The network updates are represented by the functions *f(*·) and *g(*·):

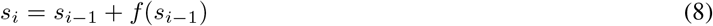

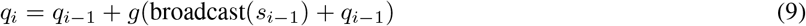

The iterative process can be organized as follows:

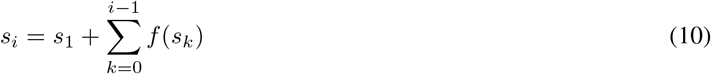

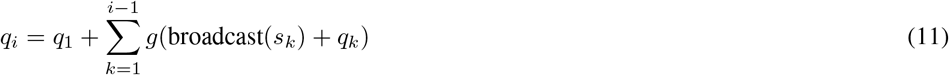

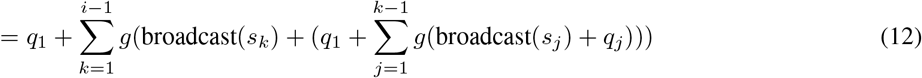

Through recursive expansion of the *q*_*i*_, we observe that each term depends on its preceding *q*_*k*_ *w*hile incorporating broadcasted features from *s*_*i*−1_ at each update step. This results in information redundancy, ultimately impeding efficient network learning and feature updating.

#### Algorithm 5

AtomAttentionDecoder

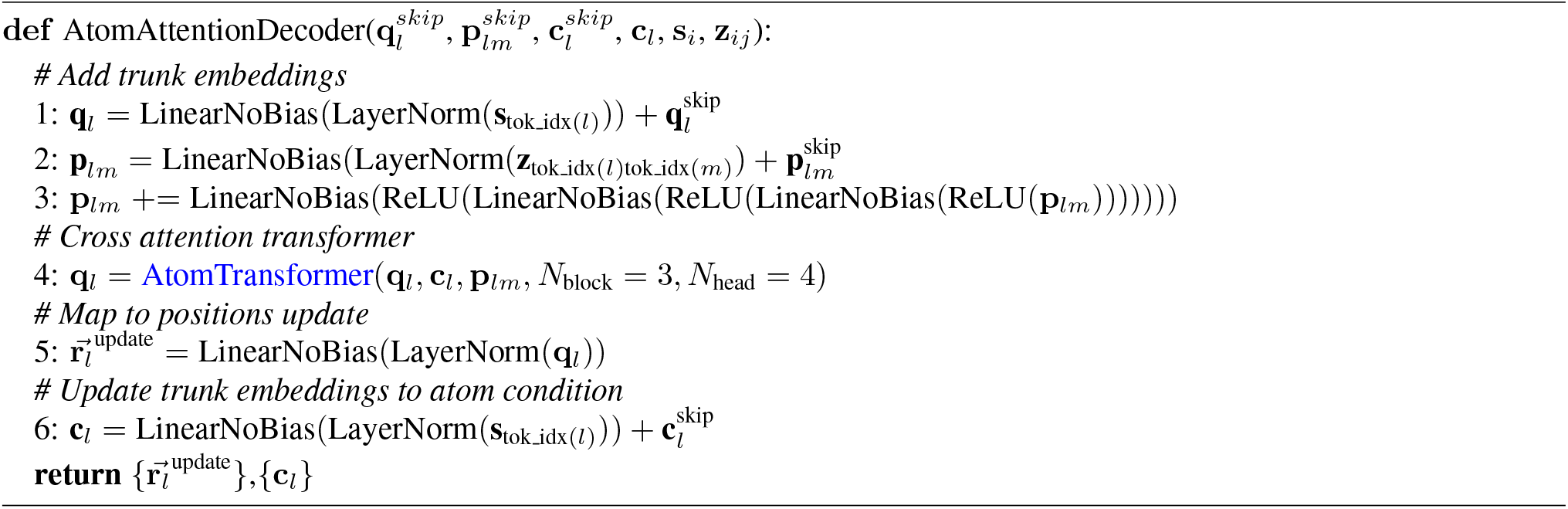

### C.6. NodeUpdate

The NodeUpdate module manages residue-level embedding updates through a time-conditioned AttentionPairBias mechanism, enabling adaptive scaling of network updates based on noise levels during the diffusion process. Additionally, we implemented dropout regularization during training to prevent overfitting and improve model generalization capabilities.

### C.7. PairUpdate

The PairUpdate module exclusively employs a modified TriangleAttention algorithm, excluding TriangleMultiplication while preserving robust generative capabilities. Our implementation utilizes recycling-derived pair representations as attention biases, enhancing the model’s ability to capture long-range interactions. To ensure pair feature space consistency, we processed recycling structures using identical binning parameters as the TemplateEmbedder module, employing 39 discrete bins spanning from 3.25 to 50.75 angstroms for comprehensive distance representation.

#### Algorithm 6

NodeUpdate

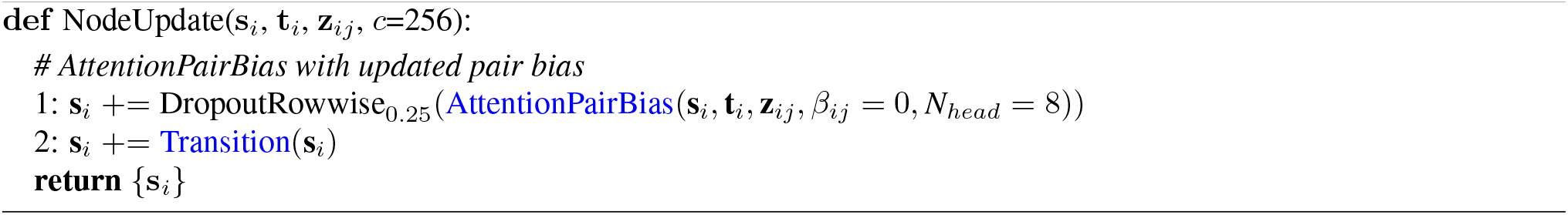

#### Algorithm 7

PairUpdate

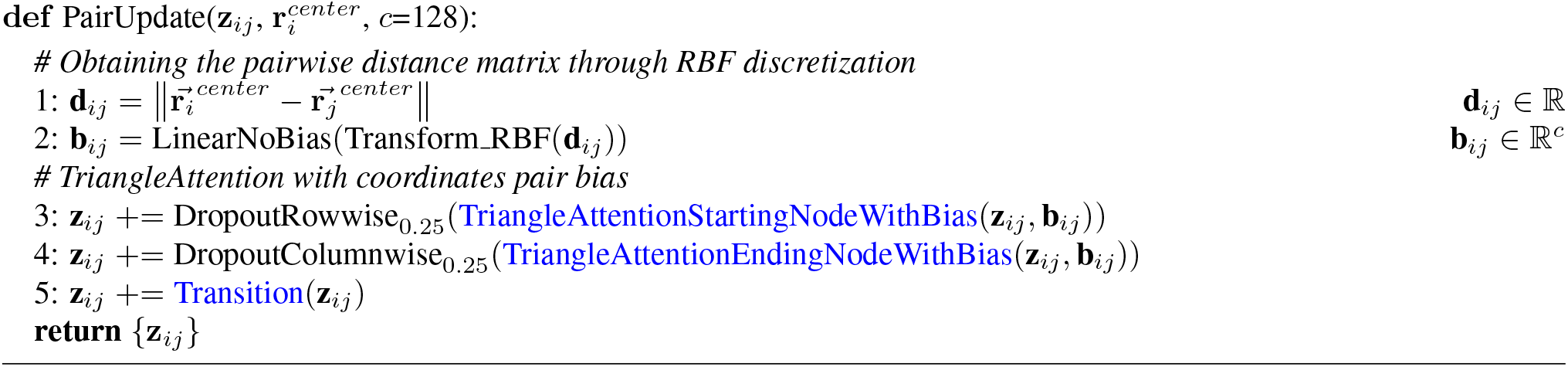

## D. Loss Function

In this paper, in addition to the primary loss function ℒ_0_, which is composed of the denoising score matching loss ℒ_*atom*_ *a*nd the cross-entropy loss ℒ_*seq*_ *f*or sequence decoding, we describe additional loss components that contribute to model optimization.

### D.1. Distogram Loss

The distogram loss function has become a fundamental component in protein structure prediction pipelines, effectively guiding the estimation of inter-residue distance matrices from pairwise feature representations. In our implementation, we symmetrize the global residue-level pair representations and project them into 64 distance bins with predicted probabilities 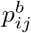, which are then supervised using one-hot encoded target distance bins 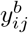.

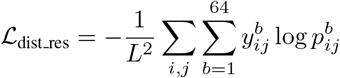

Similarly, to supervise the local relative distance distribution, we project the atomic-level pair representation from local atomic attention into 22 distance bins from 0 *Å* to 10 *Å* with 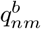, and construct the loss function using one-hot encoded target distance bins, with the local region defined by the attention window within the the 14*L* ×14*L a*tomic-level map, ensuring precise local geometry constraints.

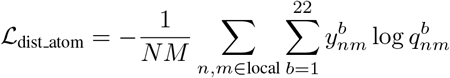

### D.2 Intermediate Loss

To stabilize the training process, we implement intermediate supervision of sequence and structure predictions from each decoder unit. Additionally, we introduce a loss weight decay mechanism with *γ =* 0.99 across decoder blocks, progressively increasing the weight assigned to later layers. Therefore, the intermediate loss for the *K u*nits can be expressed as:

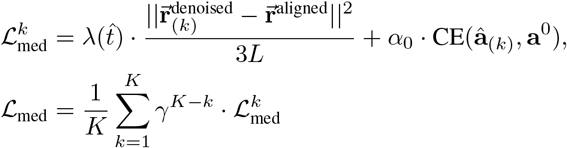

### D.3. Smooth LDDT loss function

During the training, we adopt the smooth LDDT loss (Algorithm 8, a simplified version from AF3) for all-atom protein design.

#### Algorithm 8

Smooth LDDT loss

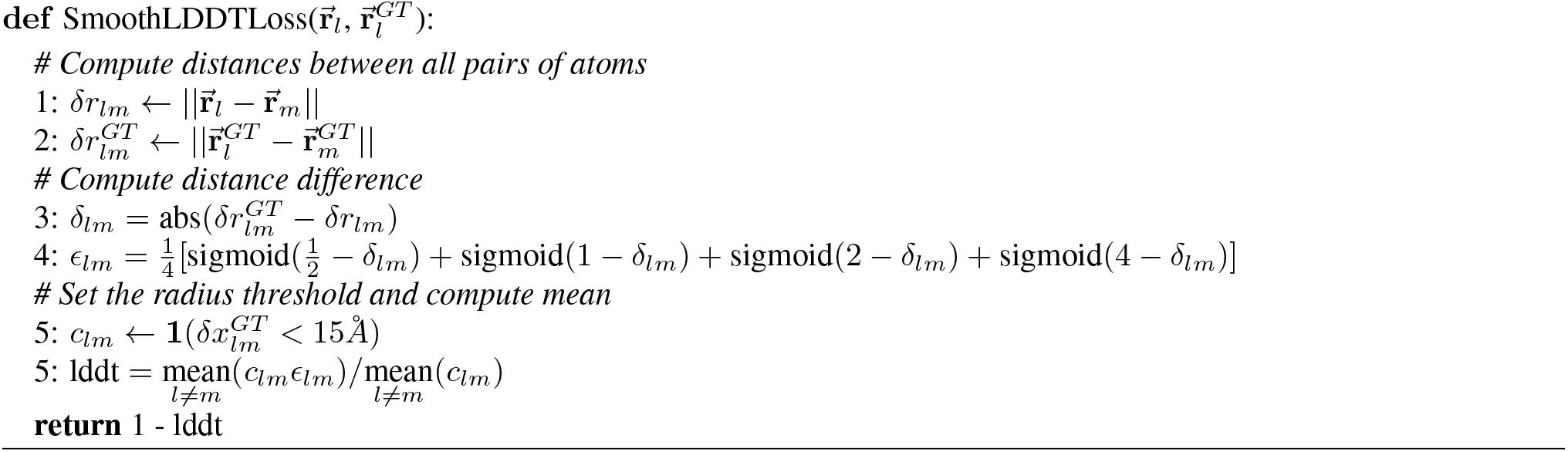

## E. Ablation Studies

We systematically investigate the fundamental factors contributing to Pallatom’s superior co-design performance through complementary top-down and bottom-up analyses, aiming to uncover the underlying principles of its exceptional designability. The success of the `atom14` all-atom representation demonstrates the viability of comprehensive atomic-level generative modeling. To further explore this paradigm, we develop a `hybrid14` representation that incorporates sequence information, enabling comparative analysis that highlights the distinct advantages of the `atom14` approach. Another key innovations in the AtomDecoder unit is the differentiable recycling process, we further investigate through ablation studies.

### E.1. hybrid14

To efficiently represent side-chain configurations across all 20 amino acids while avoiding the computational overhead of Protpardelle’s `atom73` representation, we propose the `hybrid14` representation maintains a superposition of 20 possible side-chain configurations, which collapse into a single hybridized residue state based on predicted sequence probabilities during the generation process.

Specifically, we implement the model’s self-conditioning mechanism by matrix-multiplying the predicted sequence probabilities *S* ∈ ℝ^*L×*20^ with standard amino acid reference conformer coordinates *V* ∈ ℝ^20*×*14*×*3^, generating hybrid reference conformations *V*_*hybrid*_ *= SV* ∈ ℝ^*L×*14*×*3^ that are subsequently integrated by atomic embedding.

Using identical loss functions and training configurations, we retrained the model and conducted ablation studies by generating 100 protein sequences of length *L =* 100, followed by comprehensive designability evaluation to assess the impact of atomic representation modifications.

### E.2. woRC: Ablation Setup without the Recycling Process

To assess the contribution of the differentiable recycling process in the AtomDecoder unit, we conducted ablation experiments by substituting the recycling pair with a zero matrix and disabling gradient backpropagation. This modification prevents the residue-level pair embedding *z*_*ij*_ from receiving updates from partially denoised structures while maintaining the triangle update module for self-conditioning during training.

### E.3. smooth LDDT loss

We also conducted ablation studies on the smooth LDDT loss function. Table 4 show the results. The training observations revealed that the LDDT loss maintained consistently low values throughout training, suggesting it may act as an implicit violation loss to mitigate side-chain atomic clashes. Ablation experiments confirm the LDDT loss’s role in enhancing the quality of Pallatom-generated all-atom proteins, particularly by preserving critical sequence-structure self-consistency (DES-aa reduction).

**Table 4.**
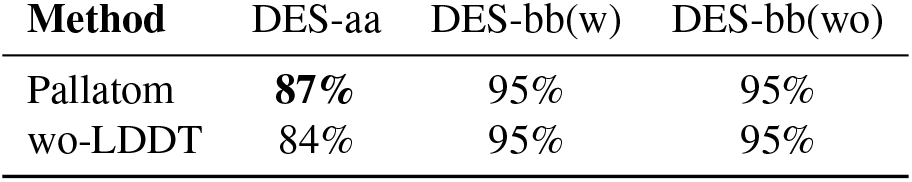
Ablation Studies on Smooth LDDT Loss.

| Method | DES-aa | DES-bb(w) | DES-bb(wo) |
| --- | --- | --- | --- |
| Pallatom | **87%** | 95% | 95% |
| wo-LDDT | 84% | 95% | 95% |

**Table 5.** Input Feature Descriptions.

| Input Feature | Dimension | Description |
| --- | --- | --- |
| ref_pos | (14, 3) | Atom positions in the reference conformer are given in Å. The backbone atoms (N, C, $C_\alpha$, O, $C_\beta$) are listed in the first five columns, while all side-chain atoms are moved to the $C_\alpha$ atom position. |
| ref_element | (14, 4) | We encode the backbone atoms based on their elemental types [N, C, O], while the side-chain atoms are encoded as a single class using ‘UNK’ (unknown). |
| ref_space_uid | ($N_{atom}$,) | Numerical encoding of the residue index associated with this reference conformer. |
| ref_center_mask | ($N_{atom}$,) | Masks indicating the center atom of the residue. |
| residue_index | ($N_{res}$,) | The pdb residue number for calculating relative positional embedding |
| residx_embedding | ($N_{res}$, 32) | The absolute position embedding by sinusoidal positional encoding. |
| template_distogram | ($N_{res}$, $N_{res}$, 39) | Pairwise distogram of pseudo $C_\beta$ are discretized into 38 bins of equal width between of bin_min=3.25Å, bin_max=50.75Å, one more bin contains any larger distances. |
| template_cb_mask | ($N_{res}$,) | Mask indicating if the $C_\beta$ atom has coordinates for the template at this residue, where 1 indicates existing residues and 0 is used for padding residues. |
| template_time | ($N_{res}$, $N_{res}$, 1) | Normalized pairwise time-step feature, ranging from 0 to 1. |
| all_atom_positions | ($N_{atom}$, 3) | The noisy position of all atoms in the system. |
| all_atom_mask | ($N_{atom}$,) | Mask indicating which atom slots are used in the the system. |
| t_hat | (1,) | The noise level value for adding noise. |

**Table 6.** Pallatom training hyperparameters.

| Parameter name | Value |
| --- | --- |
| Batch size | 32 |
| Learning rate | 0.001, No warm-up. |
| Examples per epoch | 35156 |
| Crop size | 128 |
| Loss weights | Sequence loss weight $\alpha_0 = 0.25$,<br>Smooth lddt loss weight $\alpha_1 = 1.0$,<br>Residue-level distogram loss weight $\alpha_2 = 0.5$,<br>Atomic-level distogram loss weight $\alpha_3 = 0.5$,<br>Intermediate loss weight $\alpha_4 = 1.0$<br>In the basic loss $\mathcal{L}_0$, weight allocation was applied to residue types, with a weight of 2.0 assigned to polar residues, and 1.0 to the others. |
| Diffusion timesteps ($N_{steps}$ or $T$) | 200 |
| Self-condition rate | 100% |
| EDM | Noise schedule $\text{lognormal}$. $\ln(\sigma) \sim \mathcal{N}(P_{mean}, P_{std}^2)$, $P_{mean} = -1.2$, $P_{std} = 1.5$, $\sigma_{data} = 16$,<br>Stochastic sampler $t_{min} = 0.01$, $t_{max} = 1.0$, noise level $\gamma_0 = 0.2$,<br>noise scale $\lambda = 1.003$, step scale $\eta = 2.25$ |
| Transformer | single representation dimension = 256, pair representation dimension=128, number of heads = 8, number of decoder units = 8 |
| Training Steps | $3 \times 10^5$ |
| Training time | $\approx 10$ days |
| Device | $4 \times \text{A6000}$ |

## F. Experiment Details

Table 5 provides a detailed list of the input feature.

Table 6 provides a detailed list of the hyperparameters used for training.

## G. Additional Results

### G.1. Out-of-distribution performance

#### Evaluation of Metrics

We performed extensive evaluation of Pallatom’s performance on out-of-distribution sequence lengths, specifically testing its generalization capabilities on proteins significantly longer than those encountered during training. Specifically, we generated and analyzed 250 protein structures for each of six distinct lengths *L =* 150, 200, 250, 300, 350, 400, providing comprehensive evaluation across a broad spectrum of sequence lengths. Notably, while our training set employed a crop size of 128, all comparative methods were trained on datasets containing proteins with lengths up to 384, highlighting a significant difference in training data complexity. Figures 4 illustrates the comparative performance across designability (DES), structural diversity (DIV-str), and structural novelty (NOV-str) metrics for various sequence lengths under both CO-DESIGN 1 and PMPNN 1 evaluation modes, providing comprehensive insights into the model’s length-dependent behavior.

**Figure 4.**
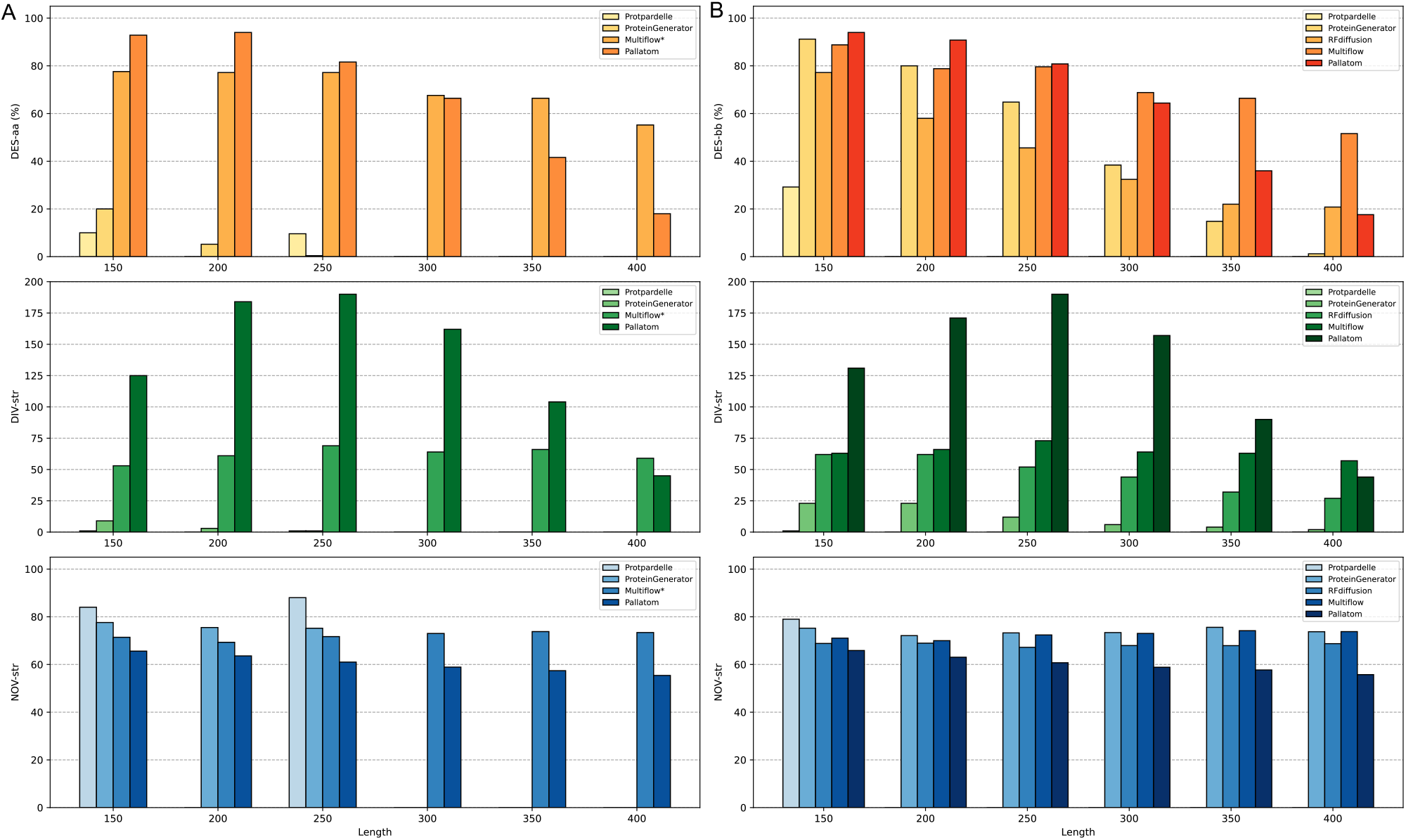
Comparison of Evaluation Metrics for Sampled Proteins at Longer Lengths. (A) and (B) respectively show the designability, structural diversity, and structural novelty under the CO-DESIGN 1 mode and the PMPNN 1 mode.

Our analysis revealed that Pallatom consistently achieved superior designability metrics for sequences up to twice the maximum training length (*L =* 150 −250) across both evaluation modes, while simultaneously demonstrating significantly greater structural diversity and novelty compared to Multiflow. Even when extending to more than three times the maximum training length at *L* = 350 and *L* = 400, Pallatom, although less advantageous in backbone design, still maintained the ability to generate all-atom proteins, a feat unachievable by the other two comparison methods, Protpardelle and ProteinGenerator.

#### Secondary Structure Analysis

We further analyzed the secondary structure preferences of sampled structures from all methods. Specifically, we utilized DSSP to classify the secondary structure of each residue in the proteins.

We established a classification scheme based on helix-to-sheet ratios: proteins with *α-*helix residues exceeding five times *β-* residues are categorized as **HHH** (all-helix), while those with *β* residues surpassing five times *α-*helix residues are classified as **EEE** (all-*β*), Proteins with balanced proportions of *α*-helices and *β*-sheets are classified as **HEL**, representing mixed *αβ* structures, completing our comprehensive secondary structure classification framework.

Figure 5 presents the secondary structure distributions for all methods under both evaluation modes, along with the corresponding distributions for designable protein subsets. This result indicates the ProteinGenerator, multiflow, and Pallatom exhibit similar secondary structure preferences within the length range of 150-400, with a roughly equal distribution between **HEL** and **HHH** structures. In contrast, RFdiffusion and Protpardelle show a stronger preference for **HEL** structures. Within the 150-400 length range, all models rarely succeed in generating **EEE** structures.

**Figure 5.**
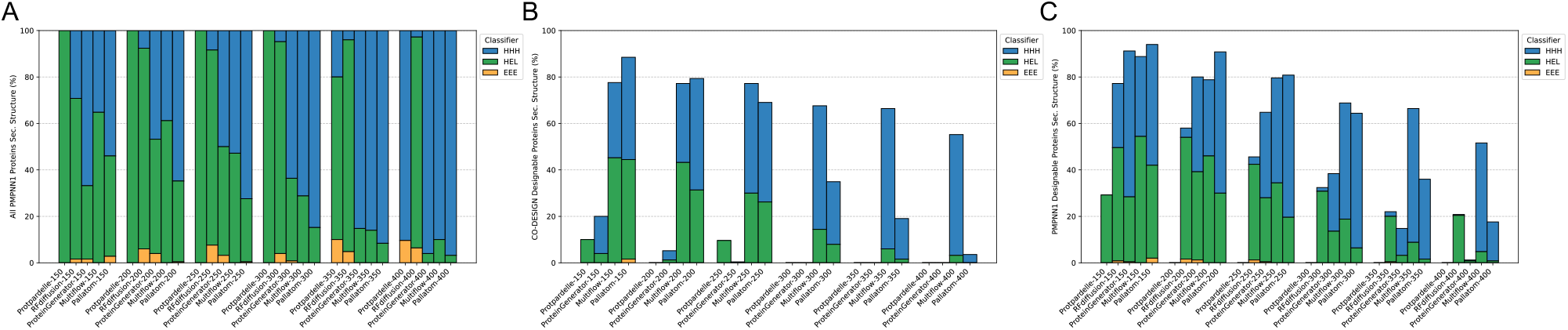
Secondary Structure Distribution of Sampled Proteins at Longer Lengths. Figures (A), (B), and (C) show the secondary structure distribution of all sampled proteins across all methods, the designable proteins in CO-DESIGN 1 mode, and the designable proteins in PMPNN 1 mode, respectively.

All these experimental results demonstrate Pallatom’s superior performance, highlighting its exceptional scalability and generalization capabilities. To further illustrate these advantages, we present additional case studies: Figure 11 showcases novel designable protein structures generated by Pallatom, while Figure 12 demonstrates its ability to produce high-quality designs for length distributions beyond the training set scope.

### G.2. Structural and Chemical Validity Assessment

As an all-atom protein generation model, we further analyzed the stereochemical properties of side chains in Pallatom-generated proteins. Through comprehensive statistical analyses of bond length distributions (Figure 6), bond angle distributions (Figure 7), *χ-*angle distributions (Figure 8), and conformational clashes (Figure 9), we demonstrate that the side-chain structures generated by Pallatom rigorously adhere to protein physicochemical constraints.

**Figure 6.**
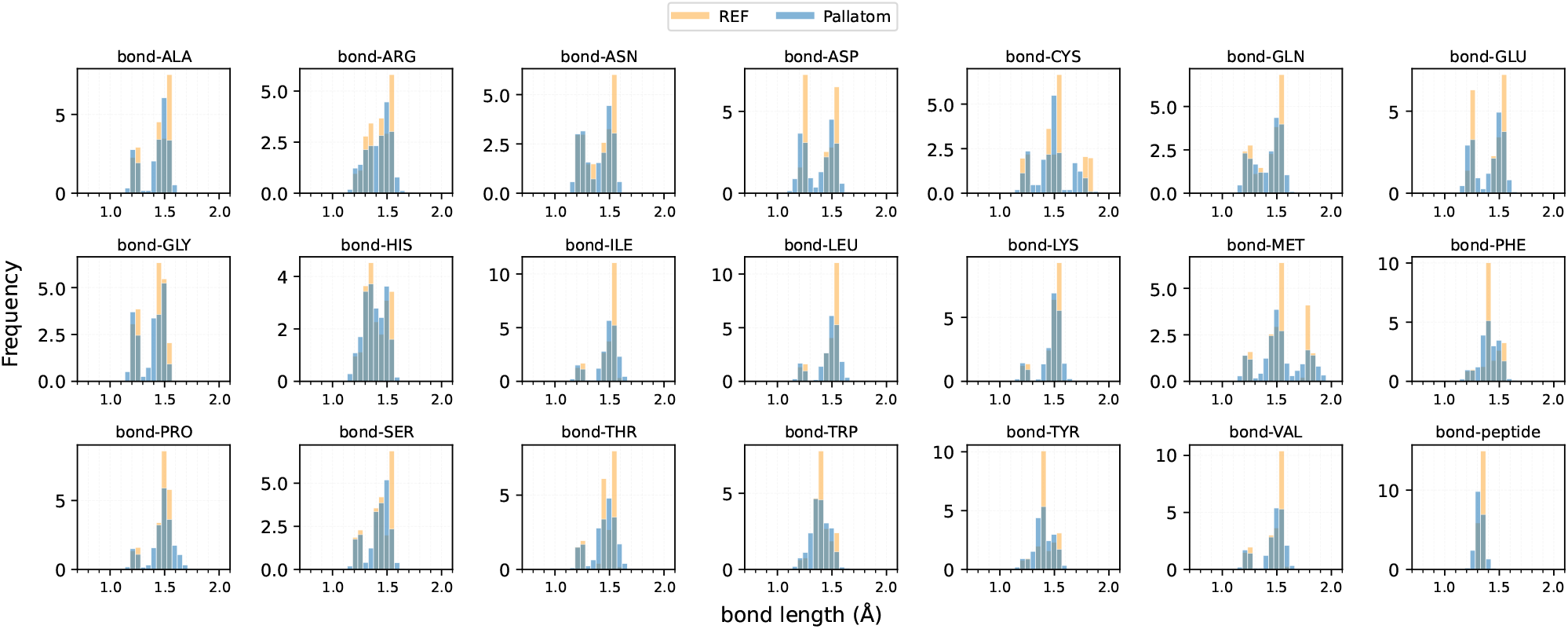
Comparison of bond length distributions between Pallatom-generated proteins (blue) and natural protein counterparts (yellow) in the PDB dataset.

**Figure 7.**
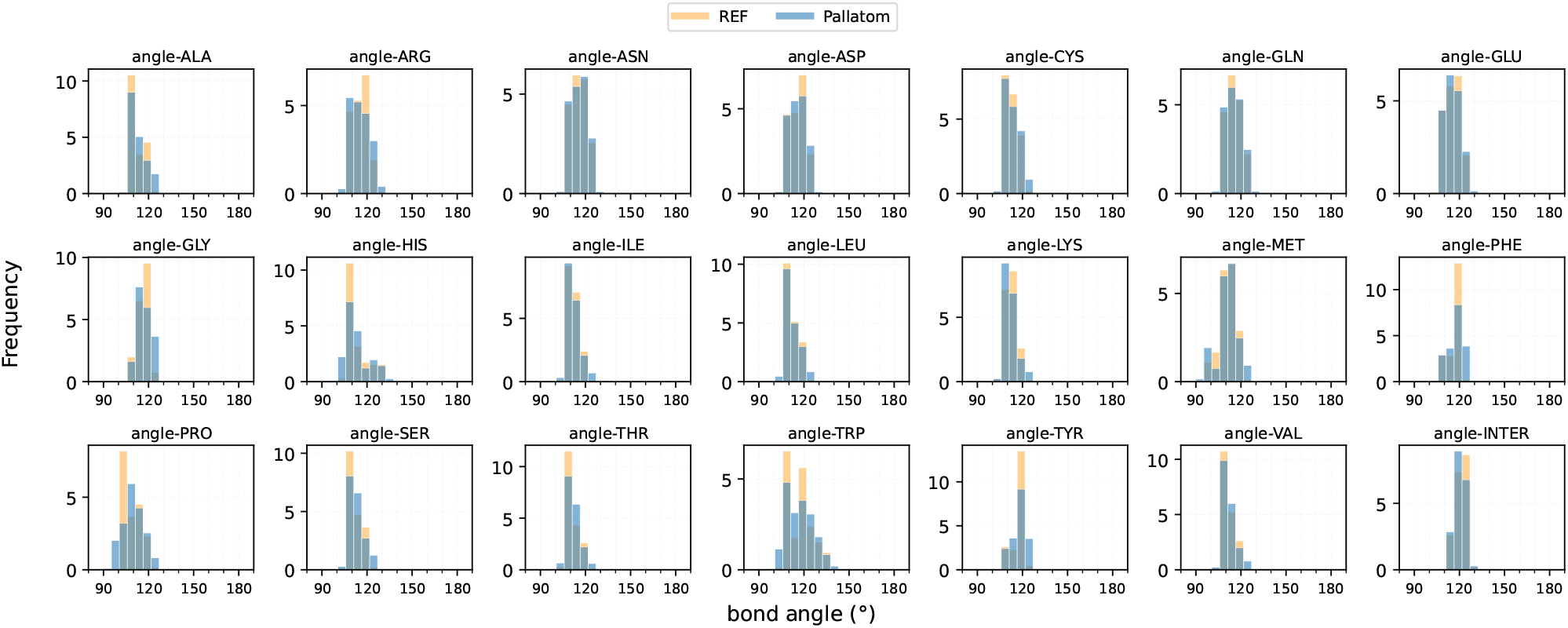
Comparison of bond angle distributions between Pallatom-generated proteins (blue) and natural protein counterparts (yellow) in the PDB dataset.

**Figure 8.**
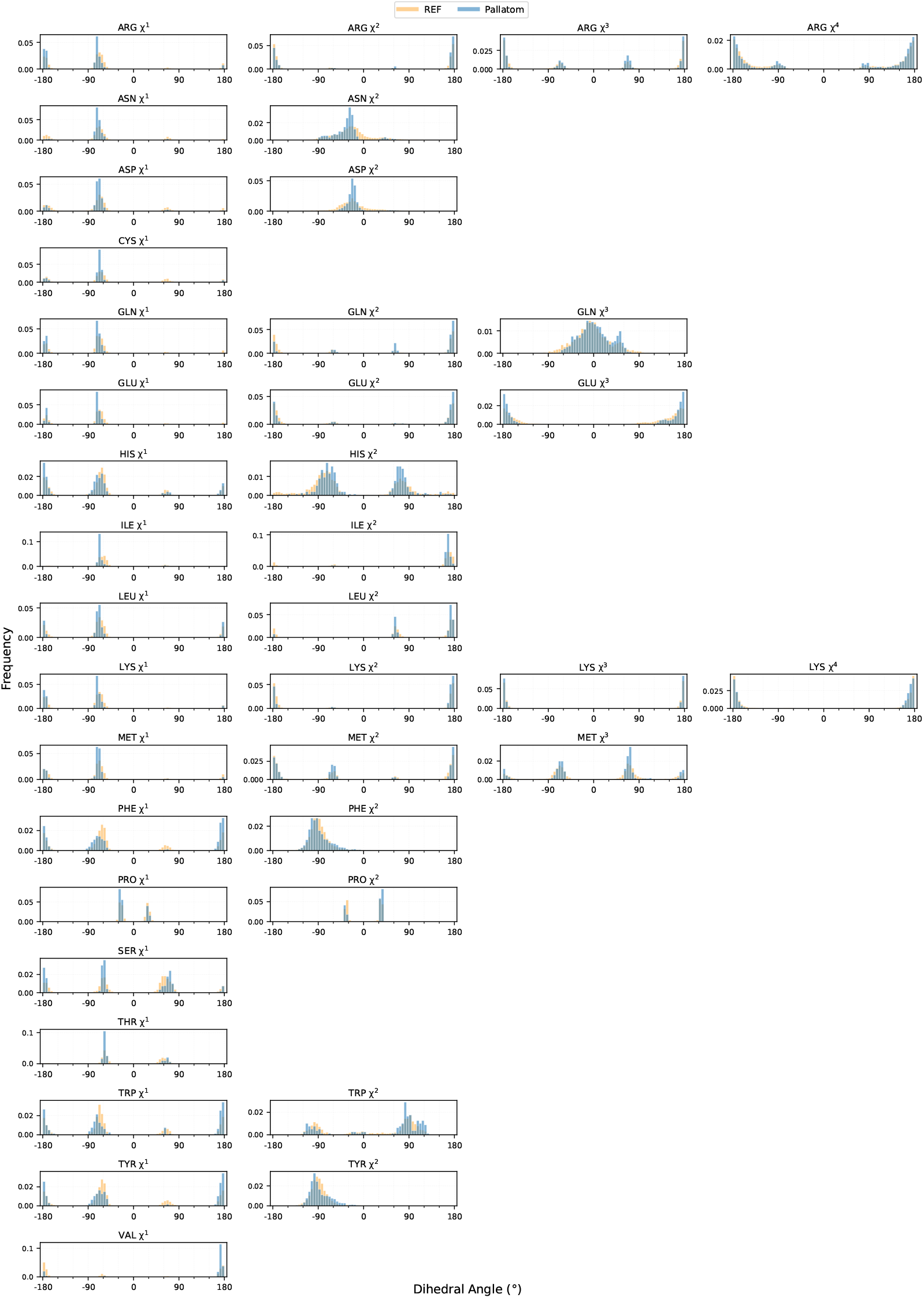
Comparison of *χ*-angle distributions between Pallatom-generated proteins (blue) and natural protein counterparts (yellow) in the PDB and AFDB dataset.

**Figure 9.**
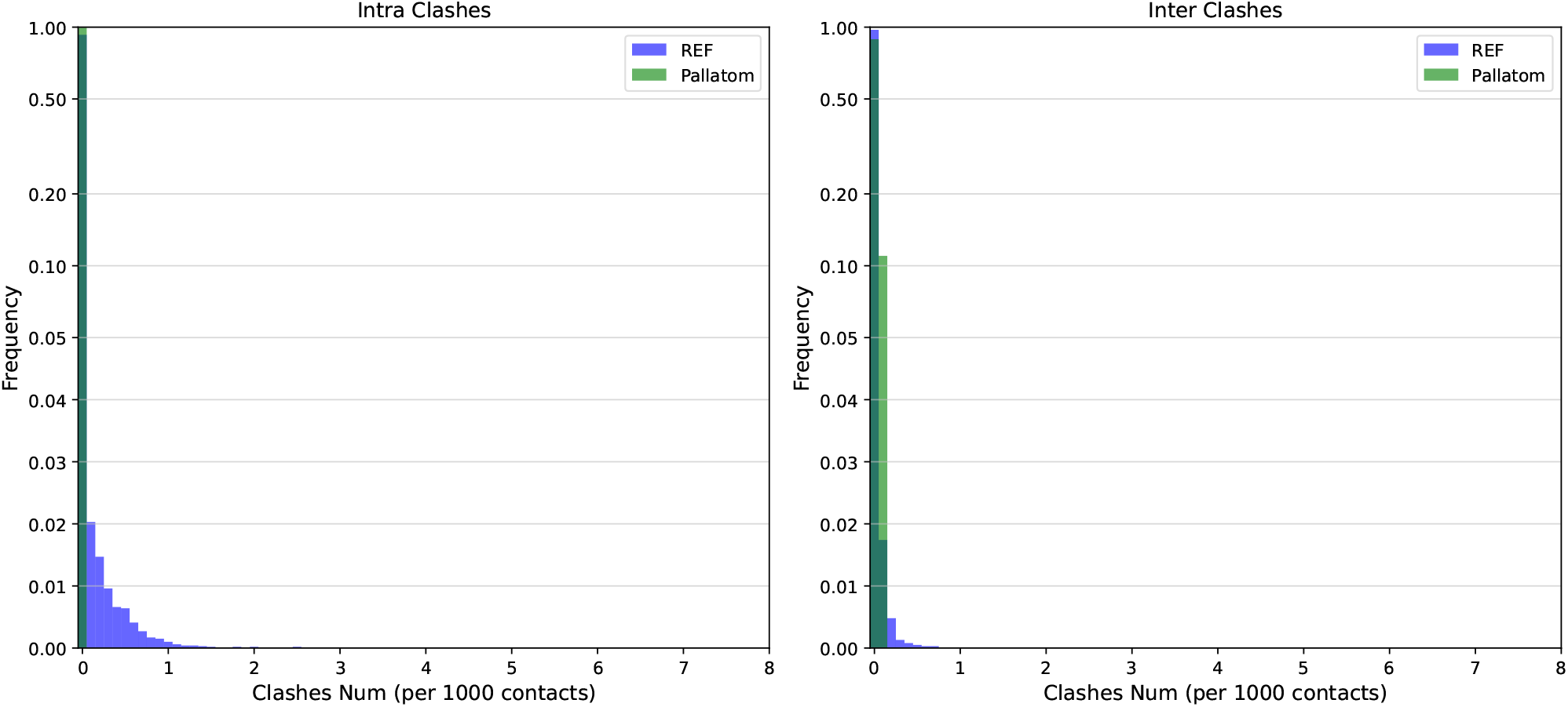
Comparison of conformational clashes distributions between Pallatom-generated proteins and natural protein counterparts in the PDB and AFDB dataset. The left panel displays intra-residue clash counts, while the right panel shows inter-residue clash counts.

Specifically, we conducted conformational plausibility evaluation on 750 generated samples (L100-L300). Conformational clashes and bond length/angle analyses were benchmarked against natural PDB datasets, while chi angle distributions were compared with training set structures (AFDB+PDB) for specific considerations: certain side chains exhibit 180° rotational symmetry where *χ a*nd *χ + π c*onfigurations are physically equivalent (differing only in atom naming) (Jumper et al., 2021). Since Pallatom was trained on both AFDB and PDB data, we used training set comparisons for *χ-*angles to avoid potential distribution discrepancies between AFDB training data and natural PDB statistics.

Bond lengths and angles were evaluated at the residue level with all-atom assessment, including separate statistics for peptide bond parameters (the peptide bond length named “bond-peptide” and the peptide bond angle named “angle-INTER”). *χ a*ngle distributions were analyzed individually for *χ*1-4 angles per residue. Conformational clashes were quantified using Clash Num (per 1000 contacts), defined as the count of non-bonded atom pairs within 5Å having distances ≤ 1.5Å ( 50% of summed van der Waals radii), categorized as intra-/inter-residue clashes.

### G.3. Analysis of Sampling Hyperparameters

We analyzed the effect of the step scale *η* on sampling proteins of unseen longer lengths. In Table 7, the left column for each length corresponds to *η* = 2.5, while the right column corresponds to *η* = 3.0. We observed that a larger step size in the gradient update direction improves the designability of proteins, and as the number of designable samples increases, the structural diversity of generated proteins also increases, with only a slight decrease in novelty. Additionally, consistent with the observations in the main text regarding the impact of sampling hyperparameters on secondary structure distribution, a larger step size is associated with a more pronounced preference for all-helix structures.

**Table 7.** Pallatom sample metrics with step scale *η* = 2.5 (left) and *η* = 3.0 (right).

| Length | 150 |  | 200 |  | 250 |  | 300 |  | 350 |  | 400 |  |
| --- | --- | --- | --- | --- | --- | --- | --- | --- | --- | --- | --- | --- |
| **DES-aa** | 88% | **93%** | 79% | **93%** | 69% | **81%** | 35% | **66%** | 19% | **41%** | 4% | **18%** |
| **DIV-str** | **131** | 125 | 153 | **184** | 149 | **190** | 73 | **162** | 38 | **104** | 9 | **45** |
| **NOV-str** | **0.650** | 0.656 | **0.618** | 0.636 | **0.594** | 0.610 | **0.576** | 0.589 | **0.551** | 0.574 | **0.525** | 0.554 |
| HHH(%) | 44% | 53% | 53% | 64% | 54% | 70% | 60% | 77% | 64% | 85% | 69% | 92% |
| HEL(%) | 53% | 44% | 47% | 36% | 46% | 30% | 40% | 23% | 36% | 15% | 31% | 8% |
| EEE(%) | 2.8% | 2.7% | 0.0% | 0.4% | 0.0% | 0.4% | 0.0% | 0.0% | 0.0% | 0.0% | 0.0% | 0.0% |

### G.4. Sequence Quality of Designable Proteins

We compared the quality of the sequences generated by Pallatom with those produced by ProteinMPNN for the same protein structures designed by Pallatom. Figure 10 shows the pLDDT scores of the two sequences predicted by ESMfold. We found that the sequence confidence score of Pallatom is slightly lower than that of ProteinMPNN, with the maximum mean pLDDT difference not exceeding 2. We attribute this difference to both the training data and the task. Regarding training data, Pallatom was trained on a monomer protein dataset, whereas ProteinMPNN was trained on a more diverse dataset that includes both monomer and multichain structures. Additionally, when preparing the training set, ProteinMPNN focused more on the sequence diversity under the same structure, while Pallatom needed to consider both sequence and structure diversity. In terms of the training task, the objectives of the two models are fundamentally different. ProteinMPNN is concerned solely with sequence design given a real backbone, whereas Pallatom must balance the dual objectives of structure generation and sequence generation from pure noise.

**Figure 10.**
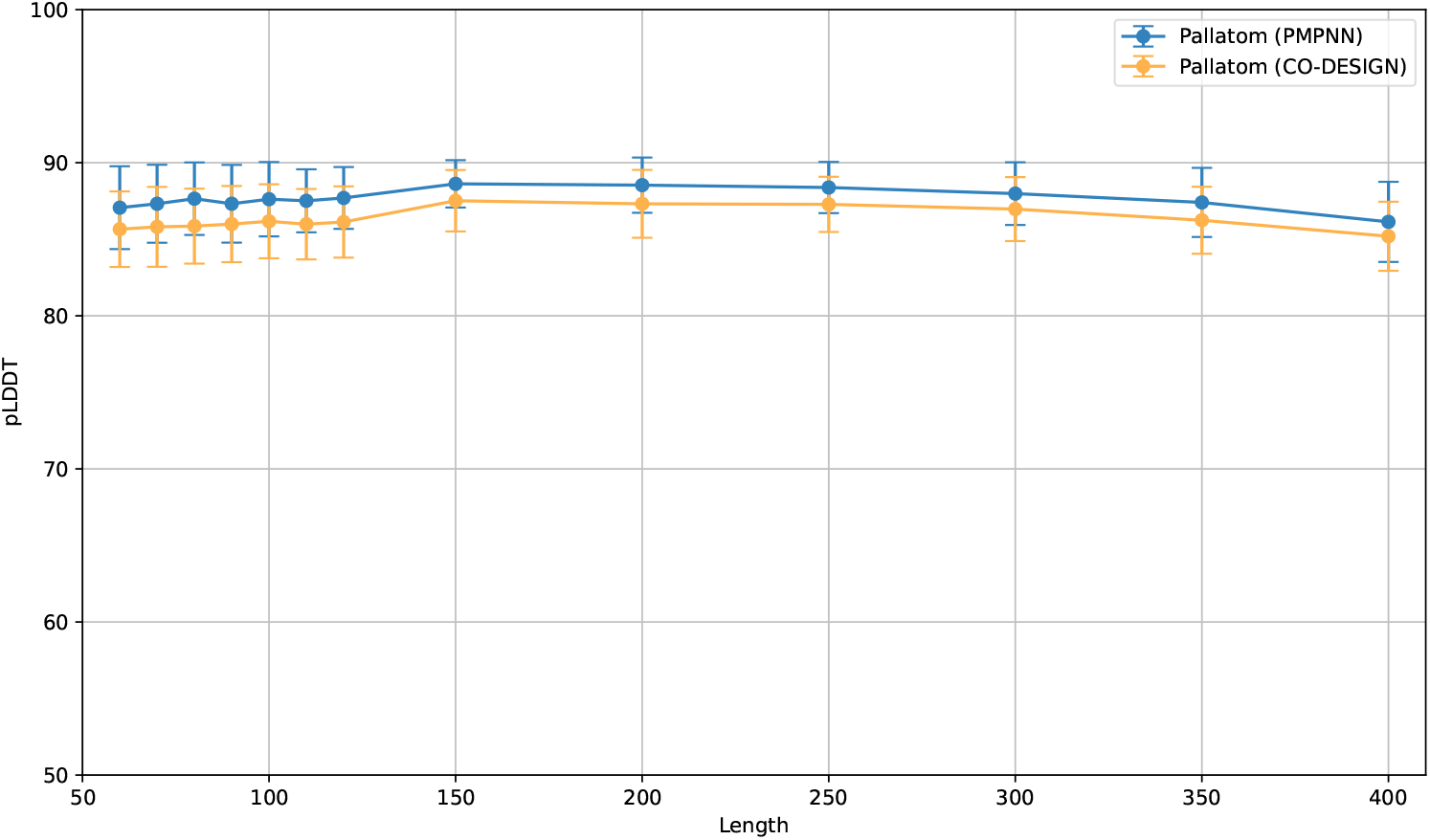
Comparison of pLDDT between sequences designed by Pallatom and ProteinMPNN across different lengths. Sequences designed by Pallatom are labeled as “Pallatom (CO-DESIGN),” while sequences designed by ProteinMPNN based on the backbone are labeled as “Pallatom (PMPNN).”

**Figure 11.**
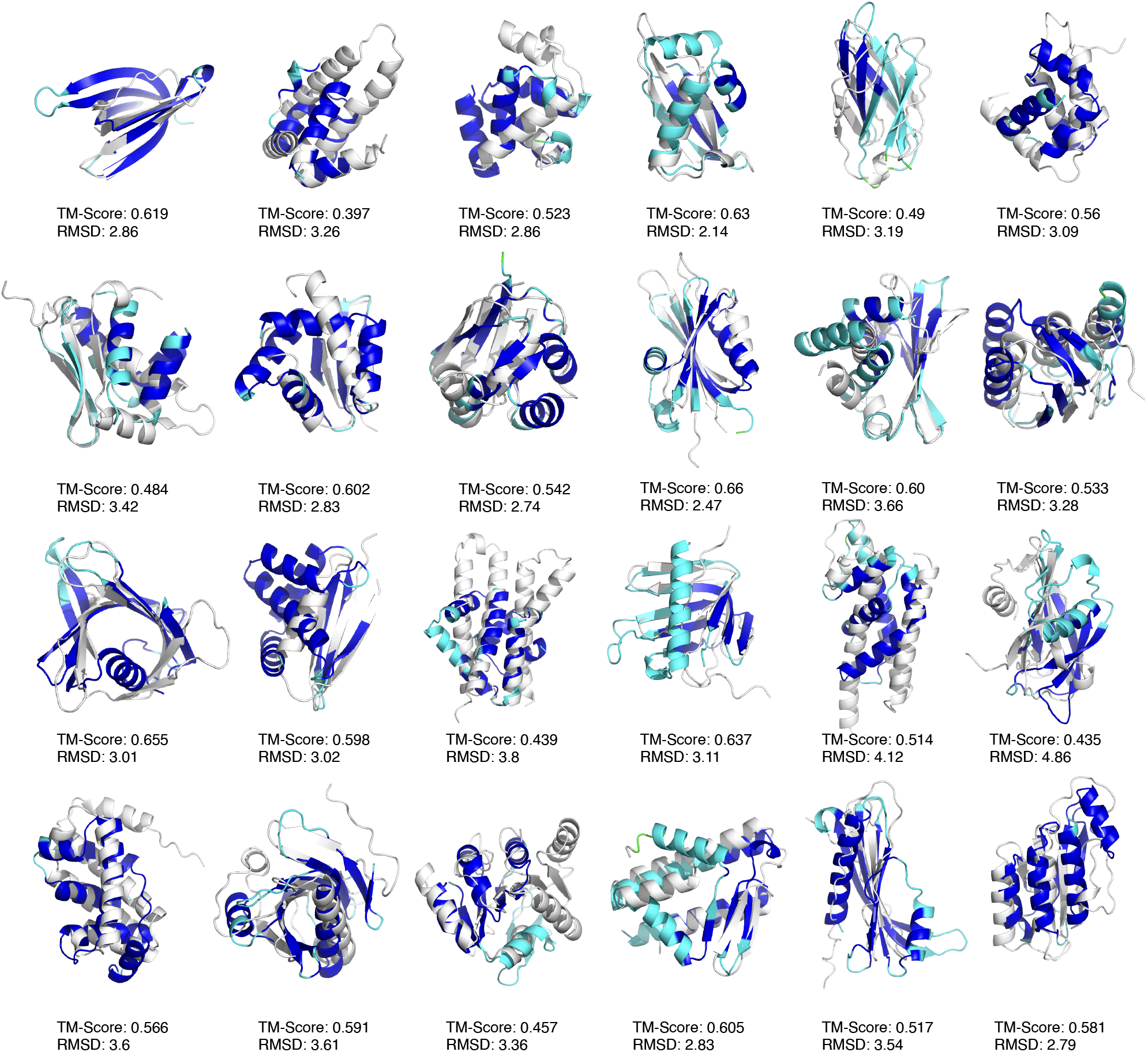
Additional novel designable proteins generated by Pallatom. The blue structures in the figure represent the designable protein sequences generated by Pallatom, which have been predicted using ESMfold and colored based on pLDDT scores. The white structures are the nearest neighbors from the Foldseek database (using the default eight databases on the Foldseek web server), with the distances between the two sets of structures evaluated using TM-Score and RMSD.

**Figure 12.**
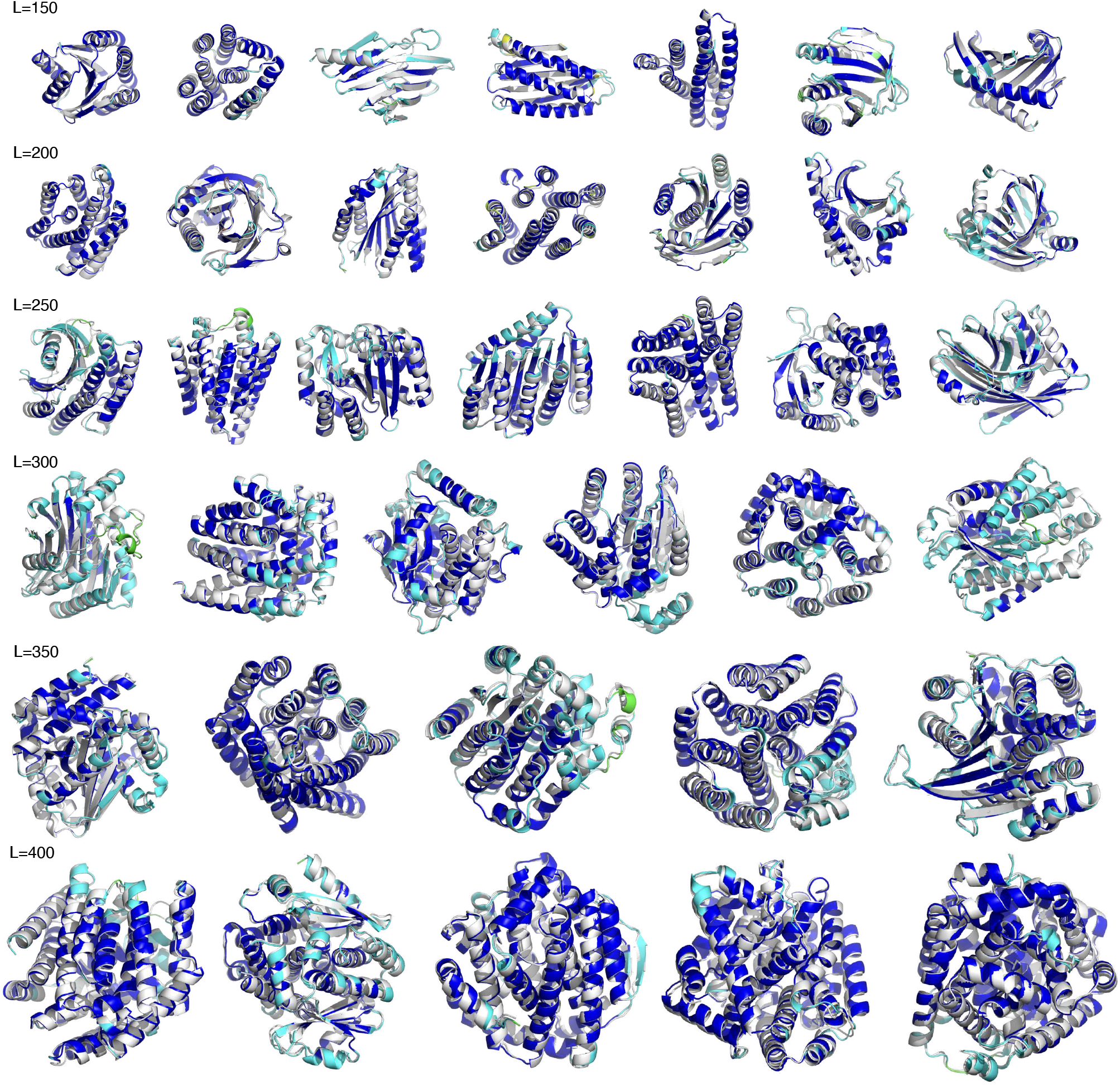
Additional all-atom protein structures generated by Pallatom for longer sequence lengths. The white structures represent those generated by Pallatom, while the colored structures are predicted by ESMfold and are colored according to pLDDT values, with bluer hues indicating higher pLDDT scores.

### G.5. Additional Comparative Methods and Results

In addition to the methodologies presented in the main text, we conducted extended benchmarking against three specialized approaches: CarbonNovo (Ren et al., 2024) (for joint backbone-sequence co-design), FoldFlow2 (Huguet et al., 2024), and Proteus (Wang et al., 2024) (both for backbone design). Systematic comparisons were performed under two modes — CO-DESIGN-1 and PMPNN-1 — with results in Table 8, 9, 10 and 11. Pallatom consistently achieved top-ranked performance across chain lengths from 100 to 300.

**Table 8.** Performance Comparison in CO-DESIGN 1 Mode.

| Metric | Method | L=100 | L=150 | L=200 | L=250 | L=300 | L=350 | L=400 |
| --- | --- | --- | --- | --- | --- | --- | --- | --- |
| DES-aa | Pallatom | **86.8%** | **92.9%** | **93.3%** | **81.0%** | **65.9%** | 41.3% | 17.9% |
|  | CarbonNovo | 75.6% | 72.8% | 70.0% | 70.0% | 59.6% | **47.2%** | **38.8%** |
| DIV-str | Pallatom | **69** | **125** | **184** | **190** | **162** | **104** | **45** |
|  | CarbonNovo | 61 | 64 | 52 | 48 | 36 | 36 | 40 |
| NOV-str | Pallatom | **0.711** | **0.656** | **0.636** | **0.610** | **0.589** | **0.574** | **0.554** |
|  | CarbonNovo | 0.717 | 0.694 | 0.717 | 0.725 | 0.748 | 0.730 | 0.701 |

**Table 9.**
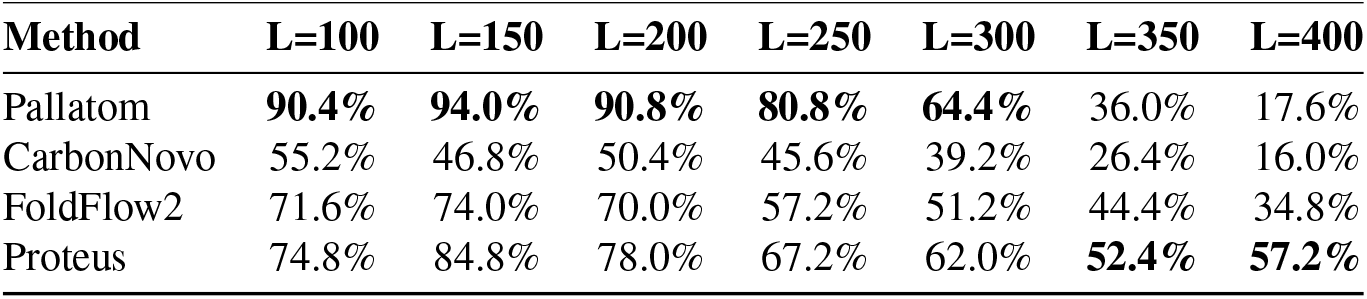
Performance Comparison in PMPNN 1 Mode for DES-bb(w).

| Method | L=100 | L=150 | L=200 | L=250 | L=300 | L=350 | L=400 |
| --- | --- | --- | --- | --- | --- | --- | --- |
| Pallatom | **90.4%** | **94.0%** | **90.8%** | **80.8%** | **64.4%** | 36.0% | 17.6% |
| CarbonNovo | 55.2% | 46.8% | 50.4% | 45.6% | 39.2% | 26.4% | 16.0% |
| FoldFlow2 | 71.6% | 74.0% | 70.0% | 57.2% | 51.2% | 44.4% | 34.8% |
| Proteus | 74.8% | 84.8% | 78.0% | 67.2% | 62.0% | **52.4%** | **57.2%** |

**Table 10.** Performance Comparison in PMPNN 1 Mode for DIV-str.

| Method | L=100 | L=150 | L=200 | L=250 | L=300 | L=350 | L=400 |
| --- | --- | --- | --- | --- | --- | --- | --- |
| **Pallatom** | **79** | **131** | **171** | **190** | **157** | **90** | **44** |
| CarbonNovo | 35 | 37 | 29 | 36 | 24 | 22 | 17 |
| FoldFlow2 | 30 | 32 | 32 | 39 | 25 | 26 | 18 |
| Proteus | 50 | 34 | 34 | 15 | 22 | 30 | 20 |

**Table 11.** Performance Comparison in PMPNN 1 Mode for NOV.

| Method | L=100 | L=150 | L=200 | L=250 | L=300 | L=350 | L=400 |
| --- | --- | --- | --- | --- | --- | --- | --- |
| Pallatom | **0.709** | **0.659** | **0.631** | **0.607** | **0.588** | **0.577** | **0.557** |
| CarbonNovo | 0.730 | 0.717 | 0.733 | 0.725 | 0.753 | 0.739 | 0.703 |
| FoldFlow2 | 0.740 | 0.694 | 0.679 | 0.656 | 0.641 | 0.636 | 0.631 |
| Proteus | 0.761 | 0.743 | 0.742 | 0.779 | 0.763 | 0.749 | 0.734 |

### G.6. Pallatom-bb: Backbone-Only Configuration

Through systematic validation, we established the viability of `atom14` as a comprehensive point cloud representation for all-atom protein coordinates, while concurrently investigating and implementing point cloud-based approaches for protein backbone description and design. To implement this, we simplified the `atom14` representation to `atom5` – a point cloud representation using four backbone atoms (N, C_*α*_, C, O) plus C_*β*_. We named this method **Pallatom-bb**. We trained Pallatom-bb using identical training data and employed PMPNN 1 for backbone design evaluation. By analogy to the right side of Table 1 and Figure 4, the experimental results for L=60 to 120 and L=150 to 400 are provided in Table 12, 13, 14 and 15.

**Table 12.** Comparative Evaluation of Pallatom-bb Under PMPNN 1 mode Across Chain Lengths 60–120.

| Method | Des-bb(w/wo) | DIV-str | NOV-str |
| --- | --- | --- | --- |
| Protpardelle | 30.00%(80.23%) | 15 | 0.747 |
| ProteinGenerator | 93.14%(96.34%) | 151 | 0.785 |
| MultiFlow | 84.69%(95.43%) | 184 | 0.744 |
| RFDiffusion | 78.29%(84.40%) | 165 | 0.805 |
| Pallatom | 89.89%(94.46%) | **318** | **0.716** |
| Pallatom-bb | **96.46%(97.49%)** | 164 | 0.760 |

**Table 13.** Comparative Evaluation of Pallatom-bb Under PMPNN 1 on Designability.

| Method | L=150 | L=200 | L=250 | L=300 | L=350 | L=400 |
| --- | --- | --- | --- | --- | --- | --- |
| RFDiffusion | 77.2% | 58.0% | 45.6% | 32.4% | 22.0% | 20.8% |
| ProteinGenerator | 91.2% | 80.0% | 64.8% | 38.4% | 14.8% | 1.2% |
| ProtPardelle | 29.2% | 0 | 0 | 0 | 0 | 0 |
| MultiFlow | 88.8% | 78.8% | 79.6% | **68.8%** | **66.4%** | **51.6%** |
| Pallatom | **94.0%** | **90.8%** | **80.8%** | 64.4% | 36.0% | 17.6% |
| Pallatom-bb | 92.0% | 80.4% | 52.8% | 43.6% | 34.4% | 36.8% |

**Table 14.**
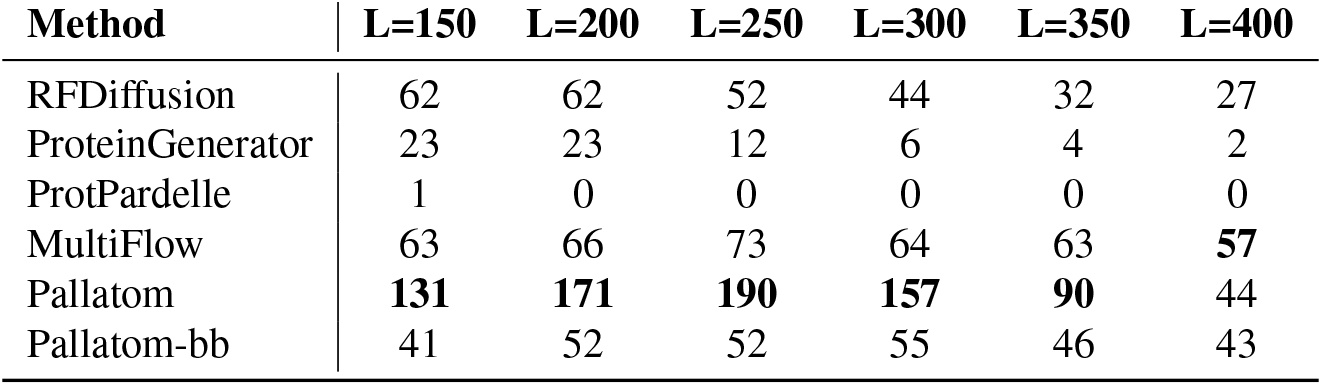
Comparative Evaluation of Pallatom-bb Under PMPNN 1 on Structure Diversity.

**Table 15.** Comparative Evaluation of Pallatom-bb Under PMPNN 1 on Structure Novelty.

| Method | L=150 | L=200 | L=250 | L=300 | L=350 | L=400 |
| --- | --- | --- | --- | --- | --- | --- |
| RFDiffusion | 0.688 | 0.689 | 0.672 | 0.679 | 0.679 | 0.687 |
| ProteinGenerator | 0.752 | 0.721 | 0.733 | 0.734 | 0.756 | 0.737 |
| ProtPardelle | 0.790 | - | - | - | - | - |
| MultiFlow | 0.710 | 0.700 | 0.724 | 0.730 | 0.742 | 0.738 |
| Pallatom | **0.659** | **0.631** | **0.607** | **0.588** | **0.577** | **0.557** |
| Pallatom-bb | 0.720 | 0.694 | 0.667 | 0.655 | 0.662 | 0.647 |

Pallatom-bb achieves high designability within training sequence lengths but exhibits reduced structural diversity. While showing limited extrapolation capability for longer sequences, it marginally outperforms RFDiffusion in overall metrics. Crucially, backbone-only point cloud approaches neglect critical side-chain/sequence interactions, whereas atom14’s all-atom framework explicitly models structural-sequence constraints, demonstrating superior holistic performance across design benchmarks.

### G.7. Analysis of Sampling Time

We conducted a comparative analysis of sampling times for each method. Specifically, we standardize the diffusion sampling steps to *T* = 200 and sample 100 proteins for each length, calculating the mean and standard deviation. All methods were tested on the same hardware: CPU: AMD EPYC 7402 @2.8GHz, GPU: NVIDIA GeForce RTX 4090 with 24GB VRAM.

Table 16 presents the results. Thanks to JAX’s JIT compilation and our optimizations at the atom level of Attention, Pallatom achieved the second fastest sampling speeds for lengths ranging from 100 to 350, outperforming all methods except Protpardelle. At *L* = 400, even with the atomic-level length reaching (14 × 400) × (14 × 400), Pallatom’s performance remains comparable to the second-fastest method, Multiflow, and is 5 times faster than RFdiffusion and 16 times faster than ProteinGenerator.

**Table 16.** Sampling Time (in seconds). The shortest time is highlighted in bold, and the second shortest is indicated in italics.

| Length | 100 | 150 | 200 | 250 | 300 | 350 | 400 |
| --- | --- | --- | --- | --- | --- | --- | --- |
| Protpardelle | *10.9±0.1* | **11.3±0.2** | **11.7±0.2** | **12.5±0.2** | **24.5±0.5** | **26.3±0.6** | **27.6±0.9** |
| ProteinGenerator | 414.1±107.5 | 389.5±2.3 | 388.8±1.3 | 477.6±1.7 | 624.0±4.1 | 796.3±4.6 | 950.3±5.3 |
| Multiflow | 25.3±0.4 | 25.3±0.6 | 27.1±0.3 | 29.4±0.2 | 35.1±0.6 | 40.5±0.2 | *46.6±0.5* |
| RFDiffusion | 95.5±14.6 | 93.9±1.0 | 106.3±0.9 | 138.5±0.8 | 183.0±0.8 | 230.7±1.6 | 287.4±6.5 |
| Pallatom* | **10.2±0.1** | *12.1±0.1* | *18.2±0.1* | *21.9±0.1* | *33.3±0.1* | *40.4±0.1* | *57.5±0.1* |

## Notes

https://github.com/levinthal/Pallatom

### Competing Interest Statement

The authors have declared no competing interest.

### Summary of Updates

Update the camera-ready version. A significant number of new diagrams, experimental results, and analyses have been supplemented in the appendix.

